# Dynamic imbalances in cell-type-specific striatal ensembles reflect learned coupling between trajectory representations and locomotor dynamics

**DOI:** 10.1101/2024.10.29.620847

**Authors:** Yuxin Tong, Brenna Fearey, Samme Xie, Andrew Alexander, Safa Bouabid, Ben Graham, Mark Howe

## Abstract

Basal ganglia models commonly propose that relative imbalances between direct and indirect pathway output shapes movement, but how such imbalances are expressed during behavior remains unclear. We simultaneously imaged identified direct-pathway and indirect-pathway spiny projection neurons (dSPNs and iSPNs) in dorsal striatum as mice locomoted through virtual visual environments for reward. Individual dSPNs and iSPNs encoded discrete locations within specific visual environments and, in a distance-based task, encoded distance traveled or elapsed time, revealing structured representations of goal-directed trajectories. At the population level, both pathways were broadly co-active and similarly correlated with locomotor speed, but their relative activity shifted systematically across learned trajectories: dSPNs dominated during early accelerating segments, and iSPNs dominated during later slowing segments. These imbalances were selectively expressed within ensembles tuned to spatial location or distance/time, depending on task structure, but were absent during comparable spontaneous locomotion outside the task context and during initial exposure to a novel environment. A computational model demonstrated that opponent plasticity driven by kinematics-linked teaching signals can reproduce the observed task-dependent imbalances through cell-type-specific plasticity of discrete trajectory-related inputs and can progressively organize locomotor kinematics over learning. Our findings indicate that direct/indirect pathway imbalances are not a general reflection of motor output, but are dynamic, state-dependent features of striatal activity that link structured trajectory representations to associated changes in behavioral vigor along repeated, goal-directed locomotor paths through learning.

## Introduction

The striatum is widely thought to regulate action through coordinated activity in the direct and indirect output pathways. A recurring feature of many influential models of basal ganglia function and dysfunction is that the relative balance of direct-pathway spiny projection neuron (dSPN) and indirect-pathway spiny projection neuron (iSPN) output is critical for regulating movement vigor, motor learning, and motor dysfunction. This idea stems from the notion that dSPNs and iSPNs oppositely regulate movement output, with dSPNs promoting movement and iSPNs suppressing it^1,2^. In classical accounts of Parkinsonian motor impairment, dopamine loss is proposed to shift this balance toward indirect-pathway output, thereby biasing basal ganglia activity toward pathological movement suppression^1,3,4^. In the healthy state, relative biases between the pathways have been proposed to support competitive interactions that regulate whether actions are initiated or terminated, as well as the vigor with which they are performed, on either moment-to-moment or learning-related timescales^5–7^. Other frameworks emphasize a role for balanced activity across the two pathways in facilitating selected actions (by dSPNs) and suppressing competing actions (by iSPNs)^8,9^.

In vivo studies have yielded a mixed picture of whether imbalances between dSPN and iSPN activity are a prominent feature of striatal dynamics. In dopamine-depleted Parkinsonian states, recordings during locomotion have revealed pathway imbalances broadly consistent with classic opponent models^10–12^. In the healthy state, signatures of imbalance have also been reported around the initiation and termination of spontaneous movements and in response to sensory cues^7,13–16^. However, other studies have shown that dSPNs and iSPNs are often co-active at similar levels at locomotion onset and can exhibit distinct but overlapping patterns of recruitment across behaviors, suggesting coordinated activity across locomotion-related ensembles^9,12,15^. Together, these findings suggest that dSPN/iSPN imbalances are not ubiquitously expressed during movement initiation, termination, or changes in vigor, but may instead emerge selectively in particular behavioral contexts in the healthy striatum.

A critical unresolved issue is what such imbalances, if present, are anchored to. Striatal activity is modulated not only by movement kinematics but also by internal and external variables, including spatial position, distance traveled, and other representations relevant to tracking task progress^17–20^. Prior work has shown that dorsal striatal activity during learned locomotor routines reflects a conjunction of contextual and kinematic variables, and that striatal activity can tile phases of learned actions, suggesting that striatal signals are organized with respect to structured task states^17,18,21^. Such signals are likely important for organizing learned action patterns that guide movement toward goals^22,23^. One clear example is locomotion along a well-practiced trajectory, in which appropriate changes in speed must be coordinated with progress toward a goal location. In this setting, relative dSPN/iSPN output may be organized not simply by kinematic parameters per se, but by task-relevant representations that track progression through learned goal-directed trajectories. Under this view, pathway imbalances may emerge selectively within neural populations representing specific states of the external world or specific stages of an ongoing locomotor sequence, rather than as uniform differences between pathways that simply mirror movement kinematics. Determining whether imbalances are tied to locomotor variables alone or whether they are mapped onto representations of behavioral progress is therefore essential for understanding how basal ganglia output is structured during natural action.

Here, we addressed these questions by simultaneously imaging identified dSPNs and iSPNs in dorsal striatum as mice traversed learned virtual locomotor trajectories for reward. We asked whether dSPN and iSPN ensemble activity remains balanced during learned locomotion or instead acquires dynamic biases across the trajectory, whether any such biases are explained by locomotion alone or are anchored to task-relevant representations of progress through the trajectory, and whether their expression depends on learning. We find that dSPN/iSPN activity is broadly co-active overall, yet exhibits systematic shifts in relative dominance across learned goal-directed trajectories. These shifts are not explained by locomotor kinematics alone, but are expressed selectively within ensembles tuned to discrete locations, times, or distances within the trajectory, and are absent during comparable spontaneous locomotion and during initial exposure to a novel environment. Together, these findings support a model in which relative pathway output reflects a learned, state-dependent bias linking representations to the locomotor dynamics appropriate for each stage of a goal-directed trajectory.

## Results

### Simultaneous imaging of dSPNs and iSPNs during locomotion through a virtual environment

We used two-photon calcium imaging to simultaneously measure dSPN and iSPN activity with cellular resolution as head-fixed mice ran through virtual environments to obtain reward. D1-tdTomato mice^24^ (n = 12) received dorsal striatal injections of AAVs driving pan-neuronal expression of the green calcium indicator GCaMP7f^25^ and were implanted with a chronic imaging window centered above the dorsal striatum^26^ (Fig. 1A). This strategy enabled simultaneous Ca^2+^ recordings from identified dSPNs (tdTomato+) and putative iSPNs (tdTomato-; Fig. 1A-B, Extended Data Fig. 1). To assess the possibility of false negatives among tdTomato-cells, we included an additional cohort of A2A-Cre mice (n = 4) injected with a Cre-dependent AAV encoding tdTomato. Recordings were collected from one to two unique imaging fields per day at locations within the dorsal medial and central striatum (Methods, Extended Data Fig. 1).

**Figure 1:**
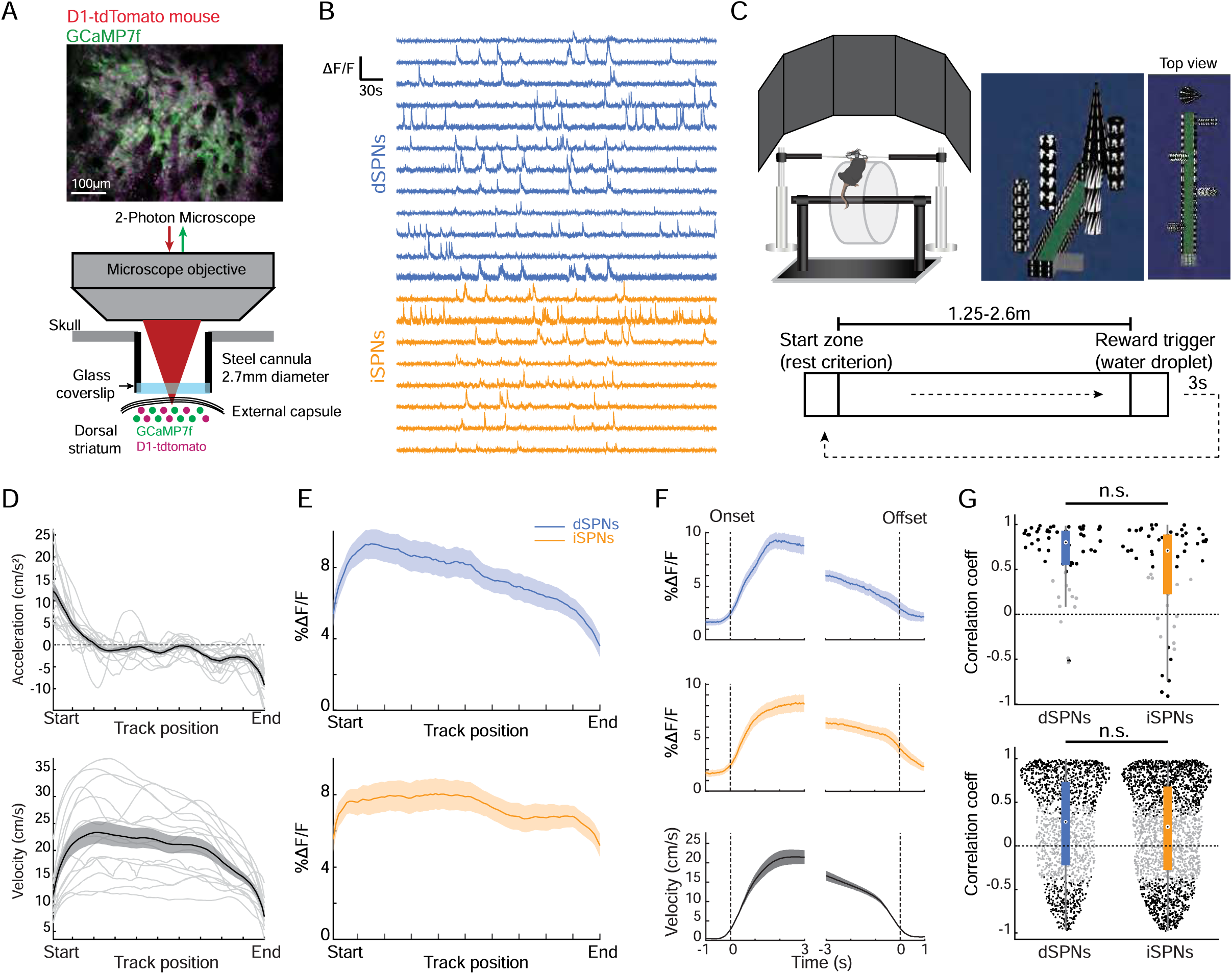
Activity of simultaneously recorded dSPNs and iSPNs is correlated with velocity during structured locomotion through a virtual linear track. **A**, Two-photon imaging approach for simultaneous calcium imaging of dSPNs and putative iSPNs in dorsal striatum. Top, representative field of view from a D1 –tdTomato mouse expressing GCaMP7f. Bottom, schematic of chronic imaging window placement over dorsal striatum. tdTomato-positive neurons were classified as dSPNs and tdTomato-negative SPNs as putative iSPNs. B, Example ΔF/F traces from simultaneously recorded dSPNs and iSPNs in a representative imaging field. C, Head-fixed virtual reality setup and linear track task. Mice ran on an axial treadmill through a virtual corridor, 1.25 to 2.6 m in length across mice, to receive water reward at the end of the track. Each trial began in a start zone after a rest criterion was satisfied, and mice were returned to the start zone after reward delivery and a 3 s consumption period. D_r_ Acceleration and velocity binned by track position during track traversal (n = 15 mice, 55 sessions). Gray traces show individual sessions and black traces indicate model-estimated means. E, Mean population ΔF/F of dSPNs (n = 1770 neurons) and iSPNs (n = 2809 neurons) binned by track position during track traversal (n = 15 mice, 55 sessions). F, Mean dSPN activity, iSPN activity, and velocity aligned to locomotor onset and offset during track traversal (n = 15 mice, 55 sessions, 58 imaging fields; 1770 dSPNs, 2809 iSPNs). Dashed vertical lines indicate onset or offset times. G, Correlations between dSPN or iSPN activity and locomotor velocity. Top, correlations between population activity binned by velocity at a fixed bin size of 0.05 cm/s and binned velocity across imaging fields (n = 15 mice, 58 imaging fields; significant fields: dSPN, 79.3%; iSPN, 75.9%; positive among significant fields: dSPN, 97.8%; iSPN, 86.4%). Bottom, velocity-binned single-neuron activity correlations with velocity (n = 1770 dSPNs, 2809 iSPNs). Black points indicate significant correlations and gray points indicate non-significant correlations. Box plots show the median and interquartile range; individual points indicate imaging fields or neurons as indicated. Shaded regions in all line plots are the 95% confidence intervals of the model coefficients from the linear mixed-effects model, n.s., non-significant comparison between dSPN and iSPN velocity correlations (p > 0.05; Wilcoxon rank sum test).

Mice were head-fixed with their limbs resting on an axial treadmill, and locomotor velocity was translated into corresponding movement through a visual virtual reality (VR) environment projected onto surrounding monitors^27^ (Fig. 1C). Animals were trained on a linear track task in which they initiated locomotion and traversed a virtual corridor containing proximal and distal visual cues to obtain a water reward delivered through a spout (Fig. 1C). Mice exhibited stereotyped locomotor trajectories, rapidly accelerating at trial onset and slowing before entering the reward zone (Fig. 1C-D; Extended Data Fig. 2A-B). These running patterns were highly consistent across trials within individual sessions, although they varied somewhat across sessions and animals (Fig. 1D; Extended Data Fig. 2A-B). Anticipatory licking was occasionally observed shortly before reward delivery but was variably present across mice (Extended Data Fig. 2C-D). Together, these behavioral features indicate that mice expressed structured patterns of locomotion along the virtual trajectory leading to reward.

We first analyzed relationships between dSPN and iSPN activity and locomotion at the population level during track traversal. Consistent with previous observations in head-fixed and freely moving rodents^17,28,29^, mean activity (ΔF/F) in both populations tracked the structured pattern of locomotor velocity across the track. Activity increased as animals initiated locomotion at the start of the track, remained elevated during sustained running, and declined as mice slowed near the end of the track (Fig. 1E-F). Mean ΔF/F during the middle portion of the track, when animals maintained high running velocity, was also significantly elevated in both populations (Fig. 1E). Overall, average population ΔF/F in both dSPNs and iSPNs was significantly positively correlated with velocity on a trial-by-trial basis, both across sessions and within individual sessions, with similar correlation magnitudes in the two cell types (Fig. 1G). Individual SPNs also exhibited significant correlations with locomotor velocity during track traversal (Fig. 1G; Extended Data Fig. 2L). Across all active SPNs, a majority of both dSPNs (56.6%; 1002/1770) and iSPNs (55.5%; 1560/2809) were significantly correlated with locomotor velocity (Spearman’s rho, p < 0.001). Of these neurons, most were positively correlated with velocity (Fig. 1G; 73.2% of correlated dSPNs and 68.7% of correlated iSPNs), consistent with the population-level correlations, whereas a smaller subset was significantly negatively correlated (26.8% of correlated dSPNs and 32.3% of correlated iSPNs). Similar velocity correlated activity at the population and single neuron levels was observed in dSPNs and iSPNs outside the virtual environment, as animals locomoted spontaneously on the treadmill in bouts comparable to track traversal (Extended Data Fig. 2E-K). Overall, these results indicate that activity in both SPN types robustly tracks locomotion velocity at population and single cell levels during both spontaneous locomotion and directed locomotion in the virtual track task.

### Dynamic dSPN/iSPN imbalances track structured locomotion along a virtual goal-directed trajectory

Next, we examined whether average population-level dSPN and iSPN activity was balanced across track traversal or whether dynamic imbalances were present. Because both pathways were recorded simultaneously within the same fields of view, we could directly quantify differences in their relative population activity across the learned trajectory. Of the total SPN population, the overall fraction of identified iSPNs was greater than that of dSPNs both during track traversal and during spontaneous locomotion outside the task context (Extended Data Fig. 3I). This difference was observed in both the D1-tdTomato and A2A-Cre cohorts, indicating that it does not arise from cell-type misclassification or from effects of static fluorophore expression. Indirect-pathway SPNs have been reported to exhibit higher intrinsic excitability^30,31^, which could contribute to this baseline difference, although its significance for net basal ganglia output remains unclear. We therefore focused on dynamic, trajectory-dependent shifts in average relative pathway activity rather than on global baseline differences between pathways.

Despite similar velocity-related activity profiles in the two pathways (Fig. 1E-G), examination of simultaneously recorded dSPN and iSPN signals revealed dynamic shifts in relative pathway balances as animals traversed the track. Average dSPN activity significantly exceeded simultaneous iSPN activity near the beginning of the track when animals initiated high-velocity locomotion, whereas average iSPN activity exceeded dSPN output near the end of the trajectory when animals slowed and terminated locomotion (Fig. 2A-B). Average activity was balanced in the middle of the track, when animals maintained high-velocity locomotion. The same pattern of imbalances was observed in the A2A-Cre cohort, indicating that it did not arise from errors in cell-type classification (Extended Data Fig. 3A-D). These effects were also present when imbalances were quantified as the relative fraction of dSPNs and iSPNs active within each position bin and after deconvolution of Ca^2+^ signals (Extended Data Fig. 4A-D and I-J). The late iSPN imbalance did not depend on anticipatory licking, as similar effects were observed on trials with and without licking (Extended Data Fig. 5D-H). Thus, although both pathways exhibit similar overall activity profiles during locomotion, their relative output shifts systematically across the learned trajectory.

**Figure 2:**
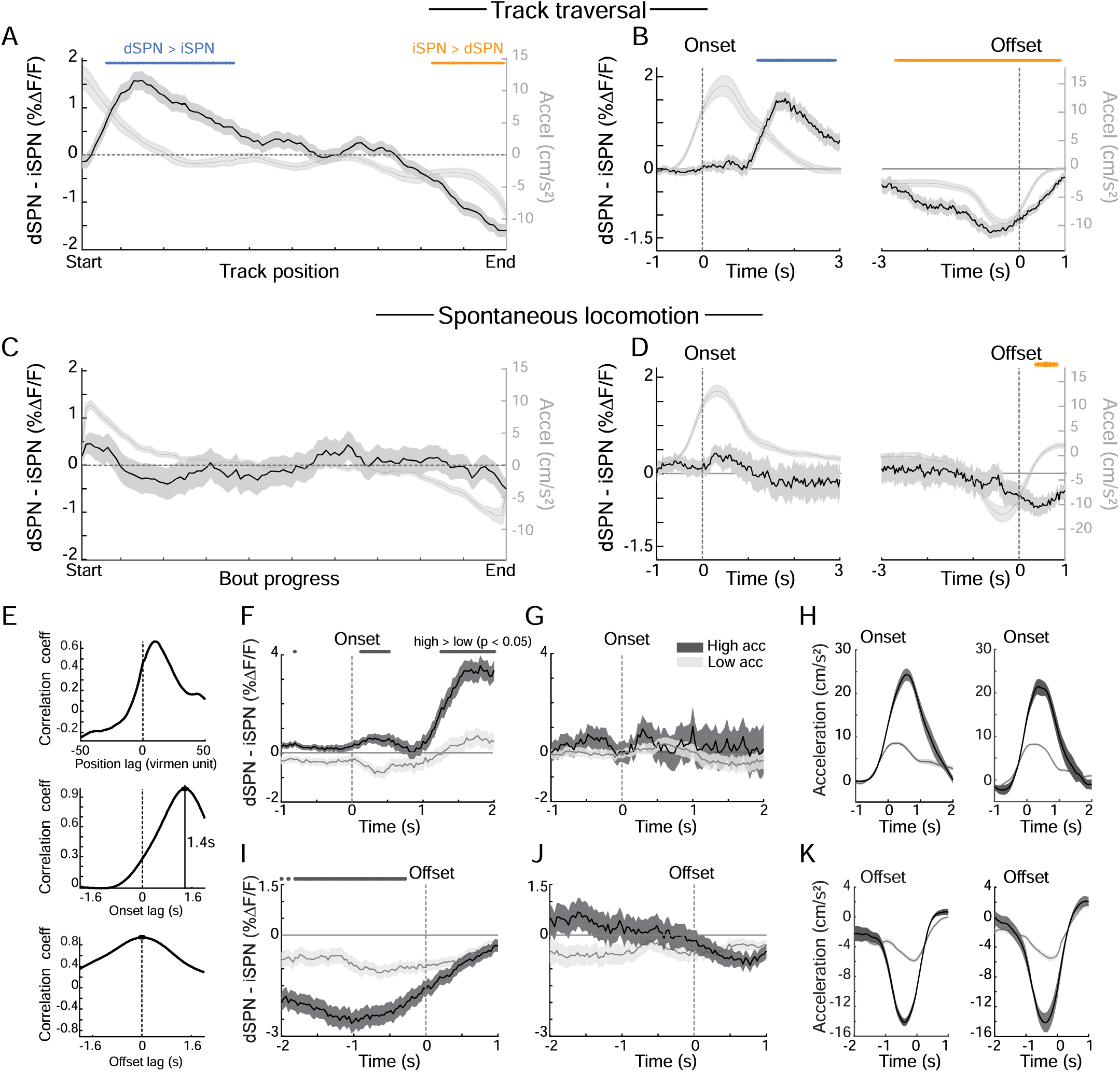
Dynamic dSPN/iSPN imbalances are expressed during structured track traversal but not during spontaneous locomotion. **A**, Mean difference in population activity between simultaneously recorded dSPNs and iSPNs binned by track position during virtual track traversal (n = 15 mice, 55 sessions, 58 imaging fields). Positive values indicate greater dSPN activity and negative values indicate greater iSPN activity. Gray trace shows model-estimated mean acceleration across track position. B_r_ Mean dSPN-iSPN activity difference aligned to locomotor onset and offset during track traversal (n = 15 mice, 55 sessions, 58 imaging fields). Gray traces show model-estimated mean acceleration aligned to the same events. Dashed vertical lines indicate onset or offset times. C, Mean dSPN-iSPN activity difference binned by normalized bout progress during spontaneous locomotion outside the structured track task (n = 9 mice, 41 sessions, 48 imaging fields). Gray trace shows model-estimated mean acceleration across bout progress. D, Mean dSPN-iSPN activity difference aligned to locomotor onset and offset during spontaneous locomotion (n = 9 mice, 41 sessions, 48 imaging fields). Gray traces show model-estimated mean acceleration aligned to the same events. E, Cross-correlation analysis between mean dSPN-iSPN imbalance and acceleration during track traversal. Top, correlation as a function of position lag. Middle, correlation as a function of onset-aligned temporal lag. Bottom, correlation as a function of offset-aligned temporal lag. Positive onset lag indicates that the dSPN/iSPN imbalance lagged acceleration. The onset-aligned correlation peaked at 1.4 s. F, dSPN-iSPN imbalance aligned to locomotor onset during track traversal for high– and low-acceleration bout groups (high group: n = 10 mice, 29 sessions, 500 onsets; low group: n = 15 mice, 47 sessions, 501 onsets; high and low groups are upper and lower tertiles based on peak onset acceleration magnitude). G, Same as F for spontaneous locomotion bouts grouped by onset acceleration magnitude (high group: n = 7 mice, 13 sessions, 44 onsets; low group: n = 8 mice, 35 sessions, 134 onsets). H, Mean acceleration profiles for the high– and low-acceleration groups shown in F and G. Left, track traversal; right, spontaneous locomotion. I, dSPN-iSPN imbalance aligned to locomotor offset during track traversal for high– and low-deceleration bout groups (high group: n = 10 mice, 39 sessions, 500 offsets; low group: n = 15 mice, 49 sessions, 502 offsets; high and low groups are upper and lower tertiles based on peak offset deceleration magnitude). **J_f_** Same as I for spontaneous locomotion bouts grouped by offset deceleration magnitude (high group: n = 9 mice, 25 sessions, 134 offsets; low group: n = 8 mice, 31 sessions, 135 offsets). K, Mean acceleration profiles for the high– and low-deceleration groups shown in I and J. Left, track traversal; right, spontaneous locomotion. Blue and orange bars indicate position bins or time points with significant dSPN > iSPN or iSPN > dSPN differences, respectively. Gray bars indicate time points where high– and low-acceleration or deceleration groups differed significantly. Shaded regions in all line plots are the 95% confidence intervals of the model coefficients from the linear mixed-effects model. Significance bars indicate p < 0.05, t-tests on model coefficients, Bonferroni corrected for multiple comparisons.

To test whether these dSPN/iSPN imbalances reflected locomotor acceleration and deceleration alone, or instead depended on the structured behavioral context of the linear track task, we examined activity during spontaneous locomotion when the VR track was turned off and rewards were delivered unpredictably. Under these conditions, mice spontaneously initiated and terminated locomotion with variable bout durations (Extended Data Fig. 2E). To compare spontaneous running with track traversal, we selected bouts with durations and peak velocities similar to those observed during VR trials (Extended Data Fig. 2E-H). Velocity encoding in both pathways was preserved during spontaneous locomotion, indicating that dSPNs and iSPNs continued to robustly track locomotor kinematics outside the task context. Thus, SPN population activity in both cell-types was broadly tuned to velocity in a context independent manner.

In contrast to the virtual track task, significant average dSPN/iSPN imbalances were largely absent during spontaneous locomotion despite similar velocity correlations (Fig. 2C,D; Extended Data Figs. 3E-H and 4E-H,M-P). No significant imbalance was detected at spontaneous bout onsets. A small iSPN-favored imbalance emerged after bout offsets, but this effect was significantly smaller than the pronounced iSPN-favored imbalance observed before track offset during VR traversal (Fig. 2D; Extended Data Fig. 5A). More generally, pathway imbalances were significantly larger during the track task than during spontaneous locomotion, both at the beginning of the trajectory, where they favored dSPNs, and near the end, where they favored iSPNs (Extended Data Fig. 5A). Together, these findings indicate that dynamic dSPN/iSPN biases are not a simple consequence of locomotor kinematics, but emerge preferentially within the structured context of the learned goal-directed task.

We next asked how these dynamic imbalances related to the structured kinematic profile expressed during track traversal. Across position bins, the average imbalance pattern was closely aligned with the acceleration profile of the learned trajectory, favoring dSPNs during the early portion of the track when animals were in an accelerating phase and favoring iSPNs during the later portion of the track when animals slowed (Fig. 2A). Activity was balanced in the middle of the track, when animals maintained high running speed, indicating that pathway biases were more closely associated with changes in locomotor state than with running speed per se. Consistent with this interpretation, the average position-binned dSPN/iSPN imbalance was strongly correlated with position-binned acceleration (Spearman’s rho = 0.69, p < 0.001, Fig. 2E). However, the dSPN-favored imbalance at the start of the track lagged the initial acceleration peak, whereas the late iSPN-favored imbalance was approximately in phase with deceleration near the end of the track (latency = 1.4s for onset and Os for offset; Fig. 2B,E). Thus, the early dSPN imbalance does not appear to reflect acceleration at the instant of locomotor initiation itself, but instead tracks, or reflects, the broader accelerating phase that occurs as animals traverse the beginning of the learned trajectory.

We next asked whether variation in acceleration magnitude was associated with variation in pathway biases, and whether differences in acceleration during track traversal and spontaneous locomotion could account for the task-dependent expression of imbalances. Trials were grouped into tertiles based on acceleration magnitude at locomotor onset or deceleration magnitude near locomotor offset. During track traversal, dSPN biased activity at onsets was greatest in the high-acceleration group and was nearly absent in the low-acceleration group, whereas iSPN dominance at offsets was greatest in the high-deceleration group (Fig. 2F,I). Because acceleration and deceleration profiles were generally consistent within individual sessions and animals, these effects likely reflect differences in the magnitude or reliability of learned kinematic structure across mice or sessions, rather than rapid trial-by-trial changes in pathway recruitment (Fig. 1D; Extended Data Fig. 2A). Importantly, spontaneous locomotor bouts with comparable high and low-acceleration or deceleration magnitudes did not show corresponding pathway imbalances (Fig. 2G,H,J,K), indicating that acceleration differences between spontaneous and track locomotion is unlikely to explain the differences in imbalances. Together, these findings indicate that dSPN/iSPN imbalances are not simple readouts of instantaneous kinematics, but are expressed in relation to phases of locomotor change within a structured trajectory through the virtual track.

### Dynamic imbalances are selectively expressed in environment sensitive, location-tuned SPN ensembles

We next asked how the population-level dSPN/iSPN imbalances were anchored to task-relevant representations in single neurons. Because the imbalances varied systematically with position along the track, we tested whether individual SPNs encoded discrete track locations. To isolate representations related to visual location, we trained a subset of mice on two visually distinct virtual tracks (n = 12 mice; Fig. 3A-B). Within each session, mice were pseudorandomly assigned to one of the two track environments on each trial. After >1 day of exposure to the second track, structured, mouse-specific velocity and acceleration profiles were highly similar between the two tracks (Fig. 3B), allowing us to isolate the influence of visual context from other variables, such as distance traveled or elapsed time. Individual dSPNs (659/1349 neurons, 48.9%) and iSPNs (1053/2650 neurons, 39.7%) displayed large Ca^2+^ transients consistently at the same track location across trials, indicating significant tuning to discrete track positions (Fig. 3C; see Methods). By contrast, other active neurons exhibited robust task-related Ca^2+^ signals but were not stably active at the same location across trials (Fig. 3C; Extended Data Fig. 6A), perhaps encoding relationships to movement kinematics, general engagement, or other un-measured task variables.

**Figure 3:**
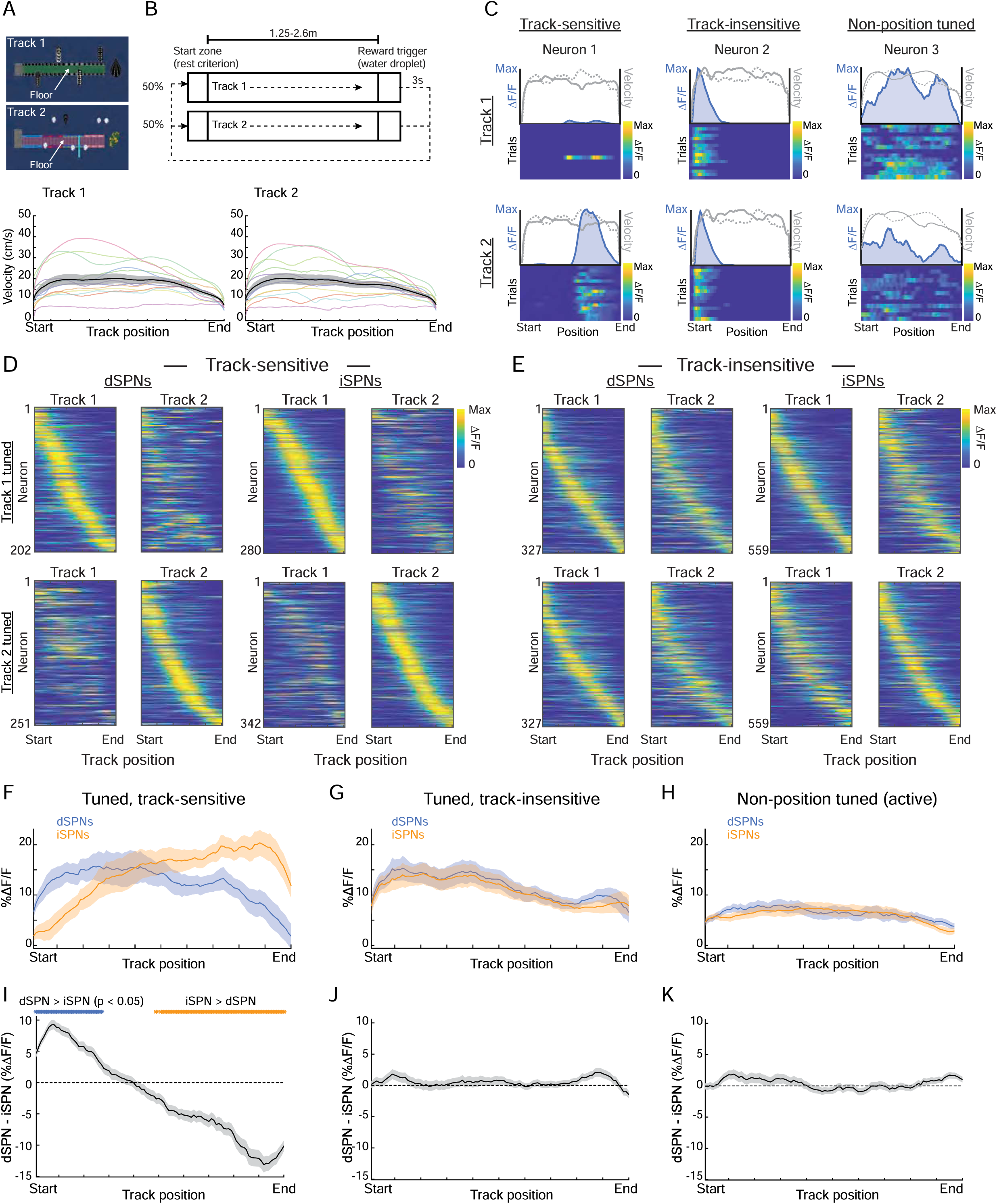
dSPN/iSPN imbalances are selectively expressed in track-sensitive, position-tuned SPN ensembles. **A**, Top-down schematics of the two visually distinct virtual tracks. **B_r_** Task schematic for sessions in which mice were pseudorandomly assigned to Track 1 or Track 2 on each trial. Bottom, velocity binned by track position for Track 1 and Track 2 (n = 12 mice, 34 sessions, 39 imaging fields). Colored traces indicate individual mice, with the same color used for each mouse across tracks; black traces indicate model-estimated means. C, Representative track-sensitive, track-insensitive, and non-position-tuned dSPNs. Line plots show mean ΔF/F for Track 1 and Track 2, with overlaid velocity traces in gray. Heatmaps show single-trial ΔF/F binned by track position. D, Population heatmaps for track-sensitive, position-tuned dSPNs and iSPNs. Top row shows neurons significantly tuned in Track 1, sorted by the position of peak activity in Track 1, and plotted for both tracks. Bottom row shows neurons significantly tuned in Track 2, sorted by the position of peak activity in Track 2, and plotted for both tracks (neurons significantly tuned to Track 1: n = 202 dSPNs, 280 iSPNs; Track 2: n = 251 dSPNs, 341 iSPNs). E, Same as D for track-insensitive, position-tuned dSPNs and iSPNs (n = 327 dSPNs, 559 iSPNs). F, Mean activity of dSPNs and iSPNs that were significantly position tuned and track sensitive in the more familiar track, plotted for Track 1 trials (n = 12 mice, 39 sessions; 202 dSPNs, 280 iSPNs). G, Mean activity of dSPNs and iSPNs that were significantly position tuned but track insensitive, plotted for Track 1 trials (n = 12 mice, 39 sessions; 327 dSPNs, 559 iSPNs). H, Mean activity of active dSPNs and iSPNs that were not significantly position tuned, plotted for Track 1 trials (n = 12 mice, 34 sessions, 39 imaging fields; 251 dSPNs, 587 iSPNs). I, Mean dSPN-iSPN activity difference for the track-sensitive population shown in F. Positive values indicate greater dSPN activity and negative values indicate greater iSPN activity. J_f_ Mean dSPN-iSPN activity difference for the track-insensitive population shown in G. K, Mean dSPN-iSPN activity difference for the non-position-tuned population shown in H. Blue and orange bars indicate position bins with significant dSPN > iSPN or iSPN > dSPN differences, respectively. Shaded regions in all line plots are the 95% confidence intervals of the model coefficients from the linear mixed-effects model. Significance bars indicate p < 0.05, t-tests on model coefficients, Bonferroni corrected for multiple comparisons.

Among significantly tuned neurons, many dSPNs and iSPNs were sensitive to the visual environment (332/659, 50.4% and 494/1053, 46.9% position-tuned dSPNs and iSPNs, respectively; see Methods), with some neurons tuned to one or two locations in only one track and others tuned to different locations in the two environments (Fig. 3C-E; Extended Data Fig. 6A-G). Other position-tuned neurons displayed similar tuning across the two tracks (position-tuned dSPNs and iSPNs, respectively), indicating relative insensitivity to visual context. At the ensemble level, both track-sensitive and track-insensitive tuned populations of dSPNs and iSPNs exhibited fields tiling the full extent of the track in both environments (Fig. 3D-E; Extended Data Fig. 6H). Field center distributions were skewed toward the beginning of the track potentially reflecting a stable, context independent representation of movement initiation (Extended Data Fig. 6H). Skewing was more prominent for track insensitive populations, indicating that stable, track invariant representations of movement initiation were more present in this population (p < 0.001 for both iSPNs and dSPNs; Kolmogorov-Smirnov test; Extended Data Fig. 6H). Together, these results indicate that both dSPNs and iSPNs contain ensembles tuned to discrete task positions, including subpopulations that are sensitive to specific visual locations within unique environments.

Next, we asked whether the dSPN/iSPN imbalances were preferentially expressed within these distinct SPN ensembles. Large, structured imbalances were present in the track-sensitive, position-tuned subpopulations (Fig. 3F,I; Extended Data Fig. 7A,D,G). Consistent with the combined population (Fig. 2A), dSPN dominance was observed during the early accelerating phase near the start of the track, whereas iSPN dominance emerged later as mice slowed near the end. In striking contrast, significantly weaker imbalances were detected in the population of position-tuned neurons that were insensitive to visual context, despite no differences in the overall activity levels in the two populations (Fig. 3G,J; Extended Data Figs. 6I and 7K). No significant imbalances were observed in the non-position tuned population (Fig. 3H,K; Extended Data Fig. 7L,M). Together, these results indicate that dynamic dSPN/iSPN imbalances arise predominantly within SPN ensembles tuned to discrete positions tiling specific visual environments as animals traverse trajectories to reach reward.

### Structured dSPN/iSPN imbalances are absent during initial exposure to a novel track

Our results indicate that dynamic dSPN/iSPN imbalances reflecting structured locomotor kinematics are organized with respect to position within a visual trajectory and are selectively expressed within subpopulations tuned to task-relevant locations. One possibility is that these imbalances emerge through experience-dependent changes in location-tuned dSPN and iSPN activity as animals acquire structured locomotor patterns along the track. Alternatively, imbalances could reflect more general features of accelerating and decelerating within a visual environment (e.g. optic flow) during reward pursuit, independent of learning.

To distinguish between these possibilities, we examined activity upon the first day of exposure to the second novel virtual environment following many days of experience in the familiar environment (Fig. 3A). Consistent with an experience-dependent emergence, dSPN/iSPN imbalances were reduced across the track in the novel environment relative to the familiar environment (Fig. 4A-B; Extended Data Fig. 8A-B). To control for differences in acceleration magnitude, we compared imbalances aligned to locomotion onset and offset for subsets of trials with matched high and low acceleration and deceleration magnitudes in the familiar and novel environments. Under these matched conditions, significant imbalances remained absent in the novel environment at both onset and offset (Fig. 4C-H; Extended Data Fig. 8C-H). Since optic flow was the same for accelerations of equal magnitude, the absence of imbalances in the novel track suggests that pathway biases depend on learned relationships between track location and structured locomotor dynamics rather than on acceleration, deceleration, or optic flow alone. Together, these results indicate that structured dSPN/iSPN imbalances are not present during initial exposure to a visual trajectory but emerge with experience as locomotor behavior becomes organized along the track.

**Figure 4:**
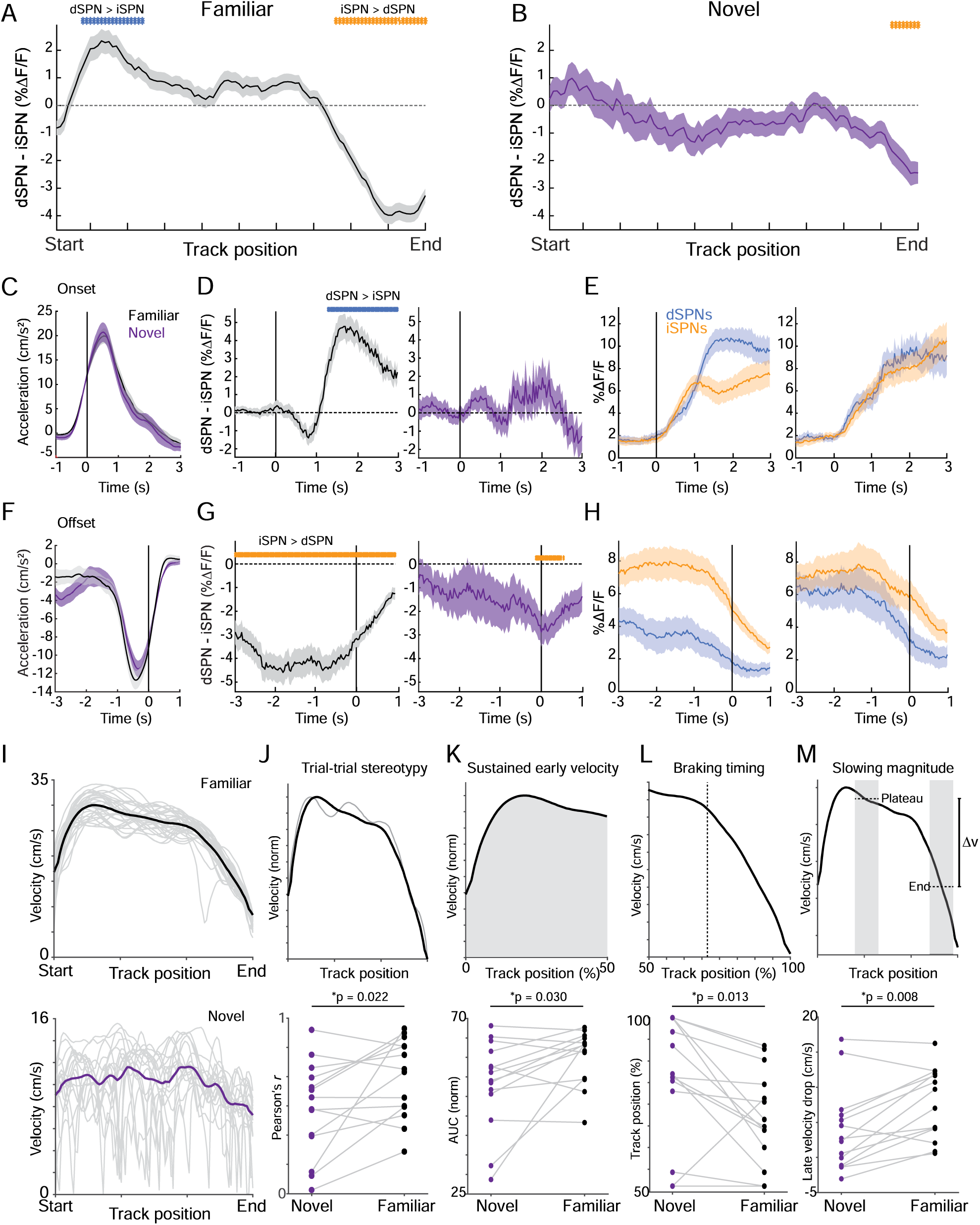
Structured dSPN/iSPN imbalances are absent during initial exposure to a novel track environment. **A**, Mean dSPN-iSPN activity difference binned by track position during traversal of a familiar virtual track (n = 14 mice, 14 sessions). Positive values indicate greater dSPN activity and negative values indicate greater iSPN activity. B_r_ Mean dSPN-iSPN activity difference binned by track position during initial exposure to a novel virtual track (n = 14 mice, 14 sessions). C, Mean acceleration aligned to locomotor onset during familiar and novel track traversal for high-acceleration onsets (familiar: n = 12 mice, 12 sessions; novel: n = 8 mice, 8 sessions). High-acceleration onsets were defined as the upper half of all onsets across mice and sessions ranked by peak acceleration. D_r_ Mean dSPN-iSPN activity difference aligned to high-acceleration locomotor onsets during familiar and novel track traversal. Dashed vertical lines indicate onset time. E, Mean dSPN and iSPN population activity aligned to high-acceleration locomotor onsets during familiar (left) and novel (right) track traversal. F, Mean acceleration aligned to locomotor offset during familiar and novel track traversal for high-deceleration offsets (familiar: n = 13 mice, 13 sessions; novel: n = 13 mice, 13 sessions). High-deceleration offsets were defined as the upper half of all offsets across mice and sessions ranked by peak deceleration magnitude. G, Mean dSPN-iSPN activity difference aligned to high-deceleration locomotor offsets during familiar and novel track traversal. Dashed vertical lines indicate offset time. H, Mean dSPN and iSPN population activity aligned to high-deceleration locomotor offsets during familiar (left) and novel (right) track traversal. I, Velocity binned by track position during familiar and novel track traversal. Gray traces show individual sessions and black or purple traces show model-estimated means. J, Trial-to-trial stereotypy of velocity profiles during familiar and novel track traversal, quantified as the mean correlation between each single-trial velocity profile and the session-averaged velocity profile (n = 14 mice; p = 0.022, Wilcoxon signed-rank test). K, Sustained early velocity during familiar and novel track traversal, quantified as area under the normalized velocity curve over the first 50% of the track (n = 14 mice; p = 0.030, Wilcoxon signed-rank test). L, Braking timing during familiar and novel track traversal, quantified as the first track position after 50% of the track where velocity fell below 90% of the plateau velocity for at least 4 consecutive bins while declining (n = 14 mice; p = 0.013, Wilcoxon signed-rank test). Plateau velocity was measured from 40 to 65 cm. M, Slowing magnitude during familiar and novel track traversal, quantified as the velocity drop from the late-track plateau window to the end-of-track window (40 to 65 cm minus 120 to 145 cm; n = 14 mice; p = 0.008, Wilcoxon signed-rank test). Blue and orange bars indicate position bins or time points with significant dSPN > iSPN or iSPN > dSPN differences, respectively. Shaded regions in all line plots are the 95% confidence intervals of the model coefficients from the linear mixed-effects model. Significance bars indicate p < 0.05, t-tests on model coefficients, Bonferroni corrected for multiple comparisons.

If these imbalances emerge through experience-dependent interactions between trajectory representations and locomotor control, then the initial absence of structured imbalances in the novel environment should be accompanied by weaker structuring of the locomotor trajectory itself. To quantify the degree to which locomotion was organized with respect to track position, we analyzed four complementary features of the position-binned velocity trajectory: trial-to-trial stereotypy, sustained early velocity, braking timing, and the magnitude of late slowing (Fig. 4I-M). Although the detailed shape and magnitude of the velocity modulation across the track varied across mice (Extended Data Fig. 8I), familiar-track behavior consistently exhibited greater spatial structure than behavior during initial exposure to the novel environment. In the novel environment, velocity trajectories were less stereotyped across trials (Fig. 4I-J), and sustained early track velocity was also reduced, consistent with weaker organization of the initial locomotor phase (Fig. 4K). Braking occurred later along the trajectory, and end-of-track slowing was diminished, indicating weaker structuring of late locomotor kinematics with respect to track position, consistent with the animals not yet linking the location of the reward with the appropriate locomotor modulation (Fig. 4L-M). Together, these results indicate that locomotion in the novel environment is more variable and less organized with respect to trajectory position, consistent with exploratory locomotor behavior and accompanied by the absence of structured dSPN/iSPN imbalances.

### Imbalances can be anchored to non location-based representations

Our results indicate that dynamic dSPN/iSPN imbalances linked to locomotor kinematics are established selectively within subpopulations sensitive to specific visual locations along learned goal-directed trajectories. In the initial track task with discrete bounded trial start and reward locations (Fig. 1C), visual location was tightly linked to reward proximity and therefore to the locomotor pattern expressed at each point along the trajectory (accelerating to a plateau velocity and slowing prior to reward). However, animals may also associate appropriate locomotor patterns with internal representations of progress through the trajectory, such as distance traveled or elapsed time. Such representations are known to exist in striatum and in upstream input regions^17,19,32–35^. We therefore asked whether visual-location associations were required for the emergence of dSPN/iSPN imbalance, or whether similar structured imbalances could also arise in a goal-directed context in which reward proximity and locomotor kinematics were linked to progress through the bout measured by distance traveled or time elapsed rather than to visual position.

To test this, we imaged dSPNs and iSPNs in mice (n = 8) trained on an “infinite track” task (Fig. 5A). In this task, mice still ran through a virtual visual environment as the previous task, but reward on each trial was delivered after they traveled a specified distance rather than reached a fixed visual location. The rewarded distance was randomly selected on each trial from a uniform distribution with mouse specific bounds (See Methods, Fig. 5A). Unlike in the bounded track task (Fig. 1C), each trial began at the location where the mouse had been rewarded on the previous trial, and the visual environment repeated every 1.5-2.2 m across mice, creating an unbounded corridor in which reward occurred at different relative visual positions across trials. Mice developed structured locomotor behavior in this task, accelerating at the start of each bout to a peak velocity and then gradually slowing as they approached the average rewarded distance (Fig. 5B; Extended Data Fig. 9G). Consistent with distance/time through the bout being more predictive of locomotor kinematics than visual location, velocity did not exhibit structured modulation as a function of visual location within the repeating track environment (Fig. 5B; Extended Data Fig. 9G). Thus, the infinite track task dissociated visual position from reward proximity and locomotor structure, allowing us to test whether dynamic dSPN/iSPN imbalances can be anchored to nonvisual trajectory representations.

**Figure 5:**
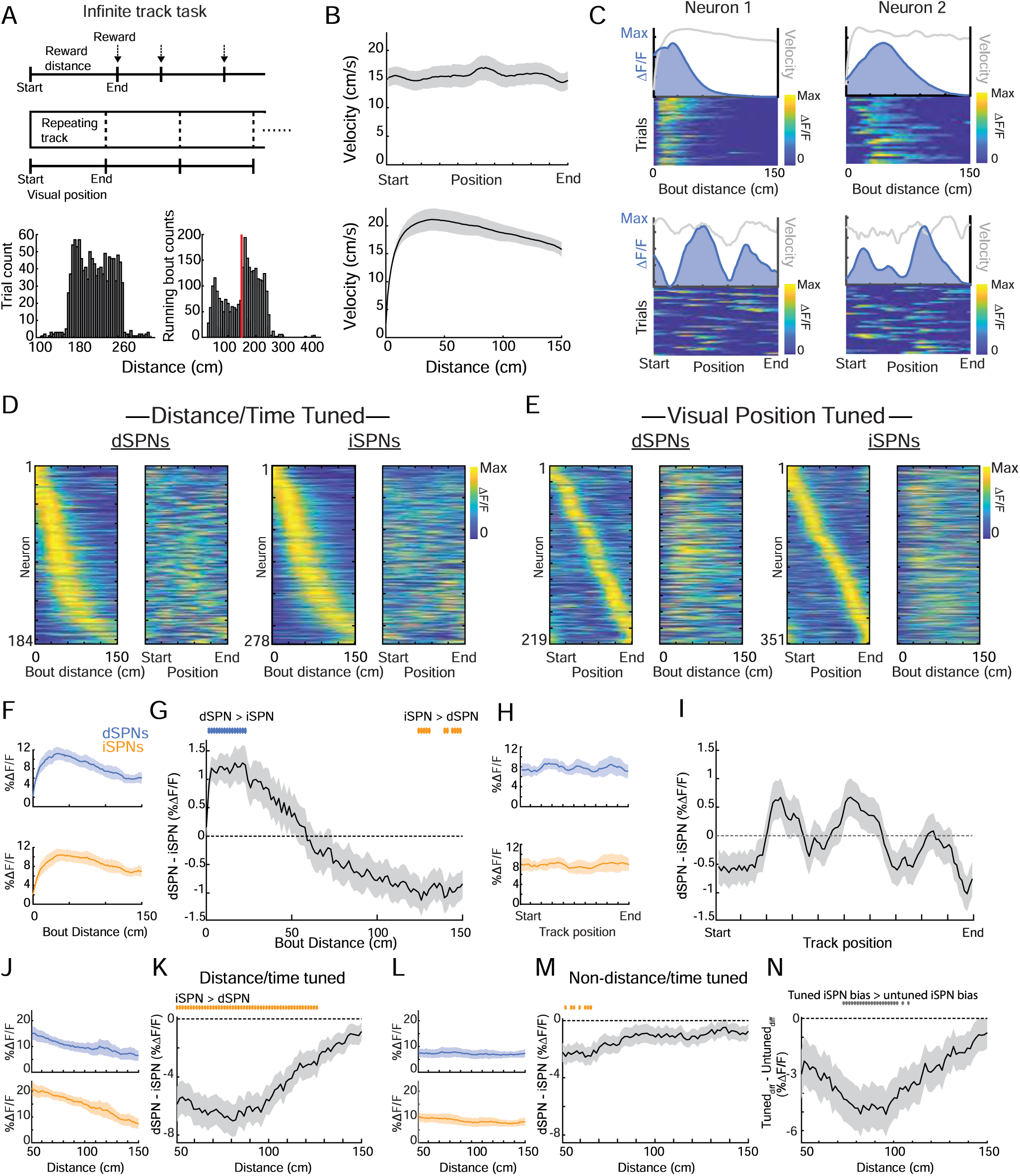
dSPN/iSPN imbalances can be organized by distance/time progress in an infinite track task. **A**, Schematic of the infinite track task. Mice ran through a repeating virtual visual environment in which reward was delivered after a randomized travel distance rather than at a fixed visual position. Top, bout distance was measured from the start of each locomotor bout to reward. Middle, visual position repeated across the unbounded track, dissociating visual position from reward proximity. Bottom, distributions of rewarded distances and running bout distances across trials (n = 8 mice, 30 sessions, 1167 rewarded trials, 1365 locomotion bouts). Red line indicates the 150 cm minimum bout-distance cutoff used to select bouts for the analyses in B to O. B_r_ Mean velocity binned by visual position within the repeating track segment (top; n = 8 mice, 30 sessions) or by distance from bout start to 150 cm (bottom; n = 7 mice, 26 sessions). C, Example distance/time-tuned dSPNs with activity binned by bout distance/time (top) or by visual position (bottom). For each neuron, line plots show mean ΔF/F and velocity, and heatmaps show single-trial ΔF/F binned by the indicated variable. D, Population heatmaps for distance/time-tuned dSPNs and iSPNs. For each cell type, the left heatmap shows activity binned by bout distance and sorted by peak distance/time field, and the right heatmap shows the same neurons binned by visual position (n = 184 dSPNs, 278 iSPNs). E_f_ Population heatmaps for visual-position-tuned dSPNs and iSPNs. For each cell type, the left heatmap shows activity binned by visual position and sorted by peak visual-position field, and the right heatmap shows the same neurons binned by bout distance (n = 219 dSPNs, 351 iSPNs). F, Mean activity of all recorded dSPNs and iSPNs binned by bout distance (n = 7 mice, 26 sessions; 611 dSPNs, 1286 iSPNs). G, Mean dSPN-iSPN activity difference for all recorded neurons binned by bout distance. Positive values indicate greater dSPN activity and negative values indicate greater iSPN activity. H, Mean activity of all recorded dSPNs and iSPNs binned by visual position (n = 8 mice, 30 sessions; 676 dSPNs, 1423 iSPNs). I, Mean dSPN-iSPN activity difference for all recorded neurons binned by visual position. **J,** Mean activity of distance/time-tuned dSPNs and iSPNs with fields after the first 50 cm of the bout, binned by distance from bout start (n = 7 mice, 26 sessions; 82 dSPNs, 132 iSPNs). **K,** Mean dSPN-iSPN activity difference for the distance/time-tuned population shown in J. L, Mean activity of non-distance/time-tuned dSPNs and iSPNs binned over the same distance range as in J (n = 8 mice, 28 sessions; 295 dSPNs, 633 iSPNs). M, Mean dSPN-iSPN activity difference for the non-distance/time-tuned population shown in L. N, Difference between the dSPN-iSPN imbalance in distance/time-tuned and non-distance/time-tuned populations, plotted across bout distance. Blue and orange bars indicate position or distance bins with significant dSPN > iSPN or iSPN > dSPN differences, respectively; gray bars indicate bins with significant differences between tuned and non-tuned imbalance traces. Shaded regions in all line plots are the 95% confidence intervals of the model coefficients from the linear mixed-effects model. Significance bars indicate p < 0.05, t-tests on model coefficients, Bonferroni corrected for multiple comparisons.

We first asked whether discrete representations of bout progress were present within SPN ensembles in the unbounded task, analogous to the position-specific representations observed in the bounded track paradigm (Fig. 3). Individual dSPNs and iSPNs were stably active across trials at particular distances from the start of each locomotor bout (Fig. 5C; Extended Data Fig. 9A-B). Since distance and elapsed time representations could not be reliably decoupled in this task, we refer to such tuning as putatively distance/time related. Overall, significant distance/time tuning was present in 184/659 dSPNs and 278/1406 iSPNs. Tuned neurons had field centers tiling the full range of bout progress analyzed, although more fields were present at earlier than later portions of the bout, likely reflecting movement initiation and/or positive velocity modulation present at the bout start (Fig. 5D; Extended Data Fig. 9B). Tuning relative to position was also present within SPN ensembles, as in the bounded track task (219/659 dSPNs and 351/1406 iSPNs; Fig. 5E). Thus, individual dSPNs and iSPNs encoded discrete distance or times within the infinite track bout when visual location was no longer predictive of reward proximity or locomotor structure.

We next asked whether structured dSPN/iSPN imbalances were expressed in the infinite track task and whether they were organized with respect to distance/time progress rather than visual position. When activity was binned by distance from the start of the locomotor bout, dSPN activity was relatively elevated early in the bout, whereas iSPN activity became dominant later as animals slowed (Fig. 5F-G). This pattern resembled the trajectory-dependent imbalance observed in the bounded track task, but was aligned to bout progress rather than fixed visual location. In contrast, when the same population activity was binned by visual position within the repeating environment, no comparable structured dSPN/iSPN imbalance was observed (Fig. 5H-I). We then asked whether the late iSPN-favored imbalance in the infinite track task was selectively expressed within neurons tuned to distance/time. Because position and distance/time tuning were partially overlapping (15.5% of dSPNs and 10.4% of iSPNs were classified as both; Extended Data Fig. 9D), we could not compare these populations directly. Instead, we compared distance/time-tuned neurons with neurons that were not detectably tuned to distance/time progress (some of these neurons were also visual-position tuned). For this analysis, we focused on neurons with fields after the first 50 cm of the bout, because activity near bout onset could reflect distance/time coding but likely also reflects movement initiation, acceleration, or velocity, similar to the track-insensitive start-related activity observed in the bounded task (Fig. 3E). Our task design could not dissociate these signals from early distance/time tuning. Using this criterion, the late iSPN-favored imbalance was prominent in second-half distance/time-tuned neurons but significantly weaker in neurons that were not tuned to distance/time progress over the same portion of the bout (Fig. 5J-N). Thus, despite overlap between visual-position and distance/time tuning, the late pathway imbalance in the infinite track task was preferentially expressed within neurons tuned to the progress variable most relevant for predicting reward. Together with the bounded track results, these findings indicate that dSPN/iSPN imbalances are not tied only to visual-location coding, but can be embedded in SPN ensembles representing time or distance within learned trajectories.

### Learned imbalances and structured locomotion can be reproduced in a computational model with opponent plasticity

Our findings suggest that dSPN/iSPN imbalances emerge when structured representations of continuous internal or external variables, such as time, distance, or spatial location, become linked to locomotor patterns reliably expressed at different stages of a repeated trajectory toward a goal. We hypothesize that such imbalances could arise through the convergence of these structured inputs with velocity-related inputs onto SPNs, and with a learning signal that bidirectionally modifies input weights according to locomotor kinematics. To formalize this idea, we developed a simplified computational model grounded in key empirical findings and prior experimental work (Fig. 6A).

**Figure 6:**
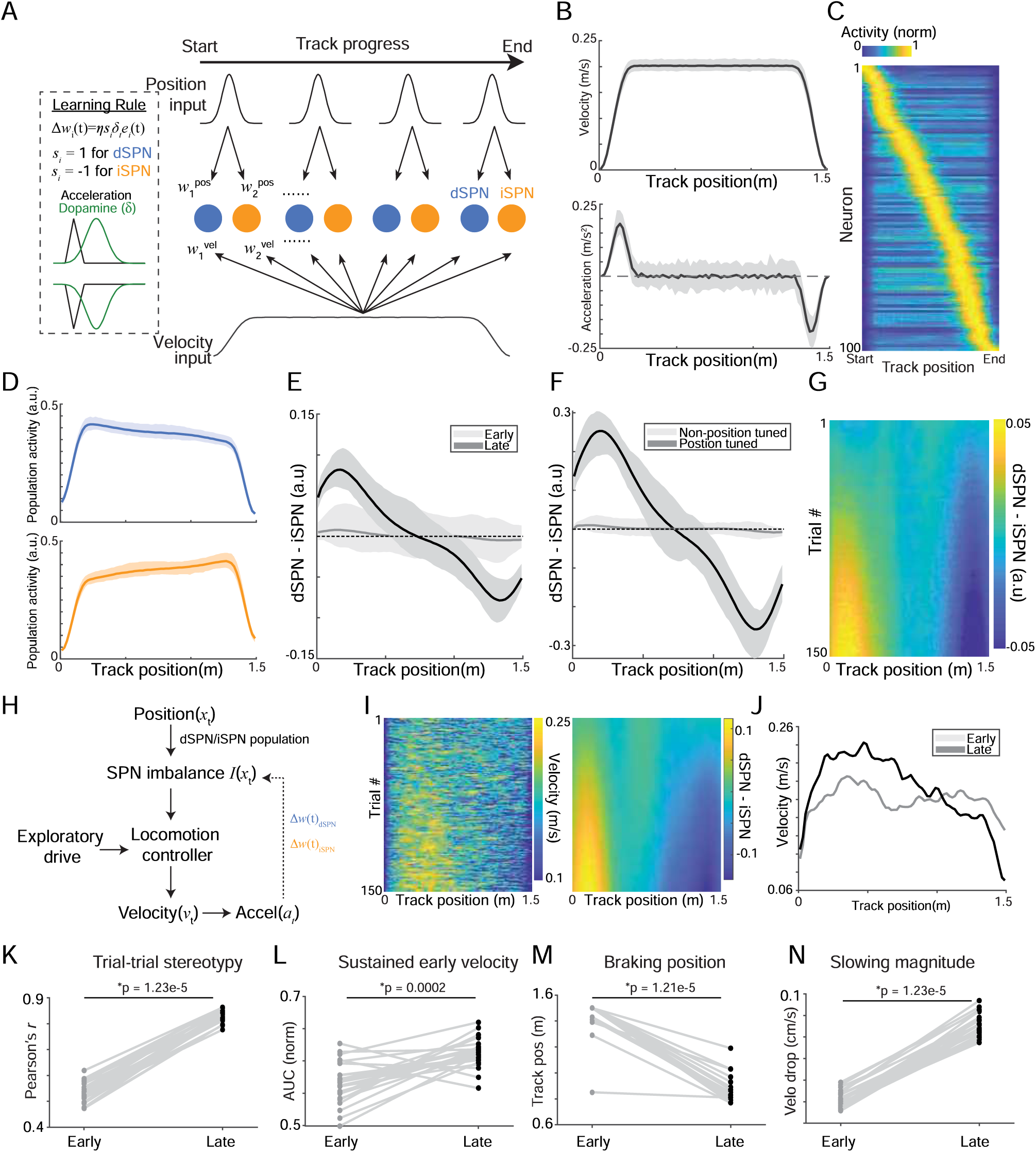
Models for the emergence of learned SPN imbalance and its contribution to structured locomotion. **A**, Schematic of the model used to examine how structured dSPN/iSPN imbalance can emerge through learning. Simulated dSPN and iSPN units received velocity input together with localized progress-related inputs tiling the trajectory. Progress-input weights were updated by an opponent plasticity rule driven by a temporally filtered kinematics-linked signal, allowing structured dSPN/iSPN imbalance to emerge over repeated traversals. B, Mean position-binned velocity (top) and acceleration (bottom) imposed in this learning model. Shaded regions indicate mean ± 95% Cl across 100 simulations. C, Example simulation showing normalized activity of the strongest progress-tuned units, sorted by the track position of peak activity. D, Mean position-binned population activity of dSPNs (top) and iSPNs (bottom) after learning. Shaded regions indicate mean ± 95% Cl across 100 simulations. E, Position-binned dSPN-iSPN difference for the first 10 versus last 10 trials of each simulation, showing the progressive emergence of structured imbalance over learning. F, The same imbalance computed separately within progress-tuned and non-progress-tuned populations, showing that the learned imbalance is concentrated in progress-tuned units. Dashed line indicates zero. G, Trial-by-trial position-binned dSPN-iSPN imbalance from a representative simulation, illustrating the progressive emergence of a structured imbalance field over repeated traversals. H, Schematic of the closed-loop model. The learned SPN imbalance feeds back onto a locomotion controller together with an exploratory drive, allowing the imbalance field established through learning to bias subsequent locomotor output. Acceleration-dependent plasticity updates dSPN and iSPN weights, linking locomotor kinematics to the progressive restructuring of behavior. I, Representative closed-loop simulation showing position-binned velocity across trials. J_f_ Position-binned mean velocity for the first 10 and last 10 trials from a representative simulation, illustrating increased sustained early velocity and earlier slowing late in learning. **K-N,** Paired summary metrics across stochastic realizations of the closed-loop model. Relative to the first 10 trials, the last 10 trials exhibited increased trial-to-trial stereotypy (K), increased sustained early velocity measured as normalized early-trajectory velocity AUC (L), earlier braking position (M), and increased slowing magnitude near the end of the trajectory (N). Each line represents one independent simulation; statistics were assessed with two-sided Wilcoxon signed-rank tests across 25 simulations.

In the model, each simulated trial consisted of traversal along a one-dimensional trajectory with a velocity profile modeled on the animals’ behavior in the linear track task: velocity increased near the start of the trajectory, remained elevated during the middle portion, and declined near the end (Fig. 6B). Each striatal unit (500 ‘dSPNs’ and 500 ‘iSPNs’) received a mixture of randomly weighted inputs encoding discrete trajectory positions and continuous locomotor velocity, and the weighted sum of these inputs determined its net activity at each time point (Fig. 6A). Position-related inputs were represented as localized fields tiling the trajectory and could be interpreted as spatial position, distance traveled, or elapsed time. Initial weights were drawn from the same distributions for dSPN and iSPN populations, so activity was approximately balanced across pathways before learning and no trajectory-dependent imbalance was built into the model. Synaptic weights on position-related inputs were then updated using a temporally filtered learning signal derived from locomotor kinematics. Specifically, this signal was a lagged, smoothed version of signed acceleration, increasing during acceleration and decreasing during deceleration. This bidirectional, kinematics-linked signal is consistent with dopamine dynamics reported in prior studies^26,36–38^ and provides a biologically plausible candidate instructive signal, although similar signals could also arise from other neuromodulatory or circuit-level mechanisms (see Discussion). Weight updates followed an opponent rule consistent with known striatal plasticity, such that increases in the learning signal strengthened dSPN-associated inputs and weakened iSPN-associated inputs, and vice versa^39–41^.

Before learning, simulated SPNs exhibited localized responses that tiled the trajectory, reflecting the structured position-related inputs, and population activity in both pathways increased with running velocity (Fig. 6C,D). Because initial weights were matched across dSPN and iSPN populations, however, average pathway activity remained approximately balanced across the trajectory and no systematic dSPN/iSPN imbalance was present (Fig. 6D,E). Through multiple trials, the model progressively developed trajectory-dependent shifts in relative pathway activity, selectively in position tuned SPNs: dSPN activity dominated early, iSPN activity dominated late, and activity remained balanced in the middle of the trajectory (Fig. 6D-G). Consistent with our data, learned imbalances were aligned with the accelerating and slowing phases of the trajectory. Thus, structured pathway imbalances similar to the experimental data emerged in our model from the interaction of structured trajectory-related inputs, kinematics-linked learning signals, and opponent plasticity.

The initial model shows that structured dSPN/iSPN imbalances can emerge when repeated locomotor patterns become associated with position-related inputs. However, during initial exposure to a novel environment, before structured imbalances were expressed, locomotion was more variable across trials and less consistently organized with respect to track position than in the familiar environment (Fig. 4I-M). This raises the possibility that imbalances may not only reflect structured locomotion but also contribute to its emergence. To formalize how this could occur, we implemented a closed-loop extension in which behavior was initially governed by an exploratory locomotion controller that produced variable velocity trajectories, with acceleration at locomotor onset and stopping in the reward zone (Fig. 6H). As dSPN/iSPN imbalances emerged over repeated traversals, they provided position-dependent feedback to the controller, such that dSPN-dominant states increased velocity and iSPN-dominant states reduced it. Under this closed-loop influence, velocity profiles became progressively more structured over learning: trial-to-trial stereotypy increased, early velocity became more sustained, braking began earlier, and late-track slowing became stronger (Fig. 6I-N). These changes recapitulated the behavioral differences observed between novel and familiar environments, in which familiar-track locomotion was more stereotyped and more strongly organized with respect to trajectory position (Fig. 4I-M). Together, these results suggest a model in which learned dSPN/iSPN imbalances arise from recurring associations between structured trajectory variables and locomotor dynamics and, with repeated experience, help crystallize stereotyped locomotor patterns along learned paths to reward.

## Discussion

Our results identify dSPN/iSPN imbalances embedded within structured representations of distance/time and spatial location along learned, goal-directed locomotor trajectories. Although dSPNs and iSPNs were similarly correlated with locomotor speed at the population level, their relative activity varied dynamically across the trajectory. dSPN activity dominated early during periods of acceleration, and iSPN activity dominated later as animals slowed. These imbalances were strongest within SPN populations tuned to the variables that defined progress through the task, including discrete locations and traveled distances or elapsed time, and were absent during comparable spontaneous locomotion and during initial exposure to a novel environment. Together, these findings argue that relative pathway imbalances are not simply a generic consequence of locomotor kinematics, but emerge with experience as striatal activity becomes linked to representations of progress through the trajectory and to the locomotor patterns associated with those states.

Classical models of basal ganglia function and dysfunction proposed that the direct and indirect pathways exert opposing influences on movement, with direct-pathway activity facilitating movement and indirect-pathway activity suppressing movement, an idea that has been especially influential in models of Parkinsonian hypokinesia^1–3,11^. However, many in vivo studies in healthy animals have reported broad co-activation of dSPNs and iSPNs during movement^9,28,42,43^. One proposed explanation that reconciles pathway opposition with co-activation is that activated dSPNs promote selected actions, while simultaneously activated iSPNs suppress other competing actions^8,44^. Our results, while not incompatible with this view, suggest an additional framework in which the two pathways are co-active in relation to the same ongoing action, but their relative balance fluctuates according to how that action is executed within a structured internal and external representations (e.g. spatial location, distance, and time) of a goal-directed trajectory. In this case, both dSPNs and iSPNs were broadly recruited during locomotion and similarly correlated with running speed, yet dSPN dominance emerged during trajectory segments associated with increasing vigor, whereas iSPN dominance emerged during segments associated with slowing or braking. These imbalances were selectively expressed in ensembles representing discrete locations, times, or distances within a trajectory and were absent during comparable spontaneous locomotion or initial exposure to a novel environment, indicating that pathway balance is not determined by locomotor kinematics alone. Instead, relative pathway dominance appears to reflect learned associations between representations of task progress and the locomotor dynamics appropriate to each stage of the trajectory. Thus, opponent biases can emerge locally within task-relevant ensembles as animals learn to structure behavior toward a goal. This perspective may also inform disease models, in which dopamine depletion could disrupt not only overall pathway balance, but also the selective, context-dependent imbalances that normally help organize learned movement.

Prior studies have reported divergent dSPN and iSPN dynamics during spontaneous locomotion in freely moving mice and across distinct behavioral motifs in the absence of an explicitly structured goal-directed trajectory^7,12,13,15^. These findings may still be compatible with our findings. One possibility is that, even under such conditions, animals develop reliable context or sensory-linked kinematic patterns that can still be understood within the present framework. In that case, the relevant task representation may not be visual position or explicit goal proximity, but another internal or external representation that aligns with progression through an unfolding behavior, such as environmental context, elapsed time, proprioceptive feedback, or progression through a motor sequence. Alternatively, some forms of pathway imbalance may arise more directly from intrinsic circuit asymmetries under particular conditions or in particular striatal regions, for example through pathway-specific input structure^45–48^ or local inhibitory interactions^49–51^.

What functional role might such structured imbalances serve? One possibility, formalized in our closed-loop model (Fig. 6H), is that they could transform initially variable, exploratory locomotion into a structured, stereotyped trajectory towards a goal. In the model, early locomotion is driven by a generalized exploratory controller that promotes acceleration and sustained movement until reward is obtained. Such a controller may arise through cortical, limbic, or other extra-basal ganglia circuits that motivate exploratory locomotion in the absence of known reward locations^52–54^. With repeated traversals, specific positions along the trajectory become consistently associated with distinct locomotor patterns: animals accelerate near the start of the track and slow near the reward zone. Kinematics-related plasticity could convert this behavioral regularity into local dSPN/iSPN biases. These biases could then feed back to influence the locomotor pattern, shaping the trajectory-specific velocity structure and promoting its consistency and automatization. This interpretation does not require the imbalances to act as instantaneous online commands for acceleration or deceleration. Consistent with this, the early dSPN bias lagged the initial acceleration peak (Fig. 2B,E). This timing suggests that pathway imbalance may reflect entry into a learned accelerating state, and may help sustain, update, or reinforce trajectory execution, rather than directly triggering locomotor acceleration. In this view, the observed imbalances may indicate a form of opponent interaction between direct and indirect-pathway circuits, consistent with work proposing competitive or opponent roles for these pathways in shaping movement vigor, reinforcement, or action control^5,6,55,56^. However, our results suggest that such opponent biases are not expressed as a global competition between action promotion and suppression. Instead, they can be anchored locally to structured representations of task progress, where they may shape how an ongoing action is executed within a learned behavioral context.

Linking pathway imbalances to representations of trajectory progress may relate to a broader role for basal ganglia circuits in the learning, stabilization, and crystallization of sequentially organized behavior. In songbirds, the basal-ganglia nucleus Area X is required for vocal learning and contributes to the shaping and maintenance of learned song structure, and lesion or perturbation studies have shown that disrupting this circuit can alter acoustic structure, tempo, and syllable sequencing^57–59^. In mammals, sensorimotor striatum has been implicated in the learning and stabilization of motor sequences, particularly in shaping their low-level kinematic structure^60–62^. Although our data address locomotor trajectories rather than action selection or discrete sequence syntax per se, they raise the possibility that learned, progress-dependent dSPN/iSPN imbalances may represent a more general striatal mechanism for stabilizing the kinematic structure of practiced behavioral sequences.

An important question raised by these findings is how structured pathway imbalances arise. Our model provides one plausible mechanism in which repeated pairings between locomotor kinematics and structured trajectory-related inputs shape dynamic imbalances across the trajectory. In this framework, locomotion-related teaching signals strengthen dSPN ensembles in trajectory states associated with increasing vigor and strengthen iSPN ensembles in states associated with slowing. In the model, plasticity was restricted to trajectory-related inputs, implemented as position-tuned fields, rather than to velocity inputs. Although this was a simplifying assumption, it captures a principle suggested by the data: pathway imbalances require an input scaffold that is consistently paired with particular kinematic phases across repeated trajectories. Velocity-related inputs are engaged during accelerations and decelerations across many contexts, including spontaneous locomotion, intertrial intervals, and home-cage running.

Bidirectional updates to these inputs would therefore be expected to average toward balance. In contrast, position (or distance/time) related inputs are active in restricted portions of a repeated behavioral context, allowing weight updates to accumulate when those inputs are reliably paired with specific locomotor dynamics. This provides one possible explanation for why structured dSPN/iSPN imbalances were largely absent during spontaneous locomotion. Another possibility is that the teaching signal itself, or the rules governing plasticity, differ between learned goal-directed locomotion and spontaneous locomotion.

What is the signal that produces bi-directional weight updates? Dopamine is an attractive candidate because it has opponent effects on dSPN and iSPN plasticity^40,41,63^. Moreover, dopamine dynamics in anterior dorsal striatum have been reported to increase during locomotor invigoration and fall below baseline during slowing, with timescales broadly consistent with the plasticity signals implemented in our model^26,36,37,64^. This is consistent with proposals that locomotion-linked dopamine dynamics could shape plasticity and progressively automate learned behavioral patterns independently of reward^65–67^. However, dopamine is not the only mechanism capable of generating structured pathway biases. Alternative or complementary contributors could include pathway-specific cortical or thalamic input structure^45–47^, local circuit interactions^49–51^ or cholinergic modulation^68,69^. Distinguishing among these possibilities will require direct measurements and manipulations of dopamine signaling and other candidate mechanisms during learning.

Several limitations should be considered when interpreting these findings. First, although our imaging approach enabled large-scale, cell-type-specific measurements of ensemble activity at a spatial scale not easily accessible with electrophysiology, calcium imaging provides an indirect and temporally filtered readout of neuronal spiking. This limits the precision with which pathway timing and output can be inferred. Although we did not detect clear timing differences between dSPN and iSPN signaling, such differences may exist below the temporal resolution of Ca^2+^ imaging and could contribute to distinct pathway roles^70–73^. Second, although our data reveal relative differences in dSPN and iSPN population activity, they do not establish how these imbalances are read out by downstream basal ganglia circuits. For example, differences in synaptic delays between the two pathways may constrain rapid-timescale integration in output structures such as the substantia nigra pars reticulata. However, multiple mechanisms within and outside the basal ganglia could extend the temporal window for integration, allowing dSPN/iSPN imbalances to influence downstream circuit dynamics or plasticity over longer timescales. Third, our imaging approach required removal of overlying cortex, primarily S1 and part of M1, both of which project to the imaged striatal region. Although mice performed the task well and expressed structured locomotor trajectories after surgery, this manipulation may have altered cortical input to the recorded SPNs. Importantly, however, several circuits that could provide structured representations of location, time, or distance, including hippocampal, entorhinal, thalamic, and other association cortical regions, were preserved, and robust locomotion-related activity was observed in both dSPNs and iSPNs^74,75^. We therefore think it is unlikely that cortical removal qualitatively accounts for the principal findings reported here. Nonetheless, future studies using less invasive approaches will be important for confirming these findings and identifying the afferent sources of progress-related striatal activity.

In summary, our findings suggest that direct and indirect pathway interactions during behavior can be understood as dynamic, learned, and state-dependent. dSPNs and iSPNs can be co-activated with similar correlates to locomotor velocity but exhibit structured shifts in relative dominance as animals progress through a learned task. These shifts are anchored to the representations of the locomotor trajectory defining progress in the current behavioral context (location or distance/time) and aligned with the locomotor dynamics associated with that progress. More broadly, our results support a framework in which the striatum links learned representations of task progress to the patterned expression of locomotor dynamics required to efficiently reach a goal.

## Methods

### Animals

Adult Drdla-tdTomato mice (Jackson Laboratory, strain no. 016204; n = 12, 9 males, 3 females; postnatal age, 12-24 weeks; body weight, 24-30 g) and wild-type C57BL/6 mice (Jackson Laboratory, strain no. 000664; *n = 4, 2* males, 2 females, postnatal age, 18 weeks) were used in this study. Mice were initially group housed and were single housed after surgery under standard laboratory conditions (20-26 °C, 30-70% humidity, reverse 12 h light/12 h dark cycle, lights on at 9:00 p.m.) with ad libitum access to food and water, except during water scheduling. During training and imaging, mice remained single housed and were maintained on a water-restriction schedule that provided approximately 1 mL of water per day. Daily water allotment was adjusted to maintain body weight at 80-90% of baseline. Of the mice used in this study, 15 were trained and imaged on the VR linear track task, 8 were imaged on the infinite track task, 9 were imaged during spontaneous running outside of VR, and 8 were imaged in both conditions. All experiments were conducted during the dark phase of the light cycle. All animal care and experimental procedures were approved by the Boston University Institutional Animal Care and Use Committee (protocol no. 201800554) and were conducted in accordance with the Guide for the Care and Use of Laboratory Animals.

### Stereotaxic virus injections and imaging window implants

Mice underwent two stereotaxic surgeries: an initial surgery for intracranial viral injection and a second surgery for chronic window implantation for two-photon imaging. For viral injections, mice were anesthetized with isoflurane (3% for induction, 1-2% for maintenance) and placed in a stereotaxic frame (Kopf). A craniotomy was made over the dorsal striatum, and AAV encoding the genetically encoded calcium indicator GCaMP7f2^5^ (Addgene; AAV1-hSyn-GCaMP7f; 3 x10^13^GC/mL, diluted 1:4 in PBS) was injected using a 33-gauge Hamilton Neuros syringe (no. 65460-02, Hamilton Company) coupled to a microsyringe pump controller (UMP3 and UMC4, WPI). A2A-Cre mice were additionally injected with AAV1-CAG-LSL-tdTomato (Addgene; 2.2 × 10^11^GC/ml). Each mouse received 300 nL of viral solution at each of four injection sites (AP, 0.4-0.6 mm; ML, 1.8-2.2 mm; DV, –1.6 to –1.8 mm) at a rate of 100 nl_/min. The craniotomy was sealed with Kwik-Sil (WPI), and the exposed skull was covered with Metabond (Parkell) to secure a metal head plate, which allowed pretraining on the linear track task (Fig. 1; see below) before window implantation.

Nine to fourteen days after viral injection, mice underwent chronic imaging window implantation^26^. Under anesthesia, the head plate was removed and the craniotomy was cleaned. The existing craniotomy was partially enlarged using a circular drill bit (FST no. 18004-27) attached to a stereotaxic drill (Foredom K.1070 Micromotor Kit). The surrounding bone was then further thinned with a handheld dental drill (Midwest Tradition 790044, Avtec Dental RMWT) and removed with forceps. Cortical tissue overlying the striatum was carefully aspirated under a surgical microscope (Leica) until corpus callosum fibers were visible. These fibers were thinned and partially removed to expose the striatal surface, but striatal tissue itself was not removed. A thin layer of Kwik-Sil was applied to the brain surface, and a chronic imaging window was inserted. Imaging windows were constructed in-house by attaching glass coverslips to metal cannulae (2.7 mm diameter^26^). Metabond was then applied around the cannula and across the exposed skull to re-secure the head plate. A metal ring centered over the window was fixed to the skull and head plate with Metabond blackened with carbon powder (Sigma). Stickers were placed inside the ring between imaging sessions to protect the window from debris.

### Two-photon imaging

Imaging was performed using a resonant-scanning two-photon microscope (Neurolabware) equipped with either a 20× objective (UMPLFLN, 0.5 NA, Olympus) or a 40x objective (LUMPIanFL N, 0.8 NA, Olympus). Excitation light was provided by an Insight X3 laser (Spectra Physics). Fields of view measured 750 × 900 μm with the 20x objective and 400 × 575 μm with the 40x objective. GCaMP7f signals were acquired at 920 nm and 31 Hz from one to two imaging fields per session, for a total of 30-70 min of imaging per day. Laser power was adjusted for each field to allow reliable detection of fluorescence transients while minimizing photobleaching. Each field was also briefly imaged at 1040 nm to visualize tdTomato expression in D1-expressing neurons. Imaging fields were sampled within an approximately 1.5 mm region centered on the cannula, spanning dorsal central-medial striatum (Extended Data Fig. 1D). To maximize the number of unique neurons sampled, the same fields were not intentionally re-imaged across days.

### Head fixed behavior apparatus and virtual track task

#### Head-fixed behavioral apparatus

Mice were head fixed with their limbs resting on a hollow 8-inch-diameter Styrofoam ball (Smoothfoam) mounted on a metal rod axle (McMaster-Carr, no. 1263K46). Ball bearings (McMaster-Carr, no. 7804K129) attached to each end of the axle were mounted in a 3D-printed cradle, allowing forward and backward locomotion while restricting angular rotation. Ball rotation was measured using an optical mouse sensor (Logitech G203; shell removed; 400 dpi sensitivity; 1 kHz polling rate) positioned parallel to the axle. Sensor output was relayed to a Raspberry Pi 3B+ and converted to an analog voltage through a digital-to-analog converter (MCP4725), then sampled at 2 kHz by a data acquisition board (NI DAQ PCIe-6343)^76^. Water rewards were delivered through a spout connected to a 60 mL syringe controlled by a digitally gated solenoid valve (Neptune Research, no. 161T012). Licking was monitored using a custom capacitive touch circuit connected to the spout. Solenoid control and lick acquisition were handled through the Nl DAQ board. Custom MATLAB software was used to control reward delivery and to visualize and acquire behavioral data.

#### Virtual linear track task

The virtual environment was built in the Virtual Reality MATLAB Engine (ViRMEn^27^) and displayed across five computer monitors arranged vertically in a semicircle, positioned approximately 35 cm from the mouse on the sides and 40 cm in front. Linear tracks consisted of distinct three-dimensional visual environments with unique distal and proximal landmarks, wall colors, and geometric patterns (Figs. 1C and 3A). Tracks were 160 virtual units in length, and input from the optical treadmill sensor was used to update virtual position based on the mouse’s locomotor velocity. The gain was scaled individually for each mouse such that track length ranged from 1.25 to 2.6 m. Custom MATLAB code was used to update virtual position and control task contingencies and trial structure.

All mice were initially trained on a 1 m track, and track length was gradually increased over training. Each trial began with the mouse positioned in a start zone, where a virtual gate blocked access to the main track. The gate opened once the mouse met a stillness criterion by maintaining velocity below 8-10 cm/s for 1.5 s. The mouse could then traverse the central arm of the track to a reward location at the end, where a 7 pL water reward was delivered with 90 or 100% probability. After reward delivery, a 5 s consumption period followed during which the display was frozen and locomotion no longer advanced virtual position. Mice were then returned to the start zone, after which a 2-6 s intertrial interval and the 1.5 s stillness criterion began. Imaging began after mice achieved a running rate of more than one trial per minute and showed clear deceleration before reward for at least 3 consecutive days.

Fifteen mice were initially trained on one track, and twelve of these mice were subsequently trained on a second environment of the same length but with distinct visual features after 5 days of experience on the first track (Fig. 3A-B). The order of environment exposure was counterbalanced across mice, such that each environment served as the familiar track in some mice and as the novel track in others. This ensured that novelty-related effects could not be attributed solely to the specific landmarks or visual features of either environment. In two-world sessions, the two virtual tracks alternated pseudorandomly across trials with equal probability. Reward probability was the same in both environments (90% or 100%). For the two-track analysis (Fig. 3), only sessions collected after at least 1 day of experience with the second track, once behavior had stabilized, were included. 15 mice were trained and imaged in VR, and 9 mice were imaged in darkness with the display turned off (Figs. 1-2 and Extended Data Fig. 2). In display-off sessions, water rewards were delivered unpredictably at intervals of 5-50 s drawn from a uniform distribution.

#### Infinite track task

In the infinite-track task, eight mice ran in a virtual environment with similar structure but distinct visual features from those used in the one-track and two-track experiments. The infinite-track was created by wrapping the mouse’s position between the start and end of a 160-virtual-unit track, without the walls defining discrete start or reward zones used in the familiar linear-track task (Fig. 1C and 5A). For each mouse, the virtual-to-physical gain was kept the same as in the familiar-track condition, such that the track length ranged from 1.5 to 2.2 m across mice. Rewards were delivered after the mouse traveled a random distance drawn from a uniform distribution where *U* ∼ (*n*一0.5, *n* + 0.5). Here *n* is a mouse specific median ranging between 1.5-2.5m that roughly matches the track length of previous experiments.

### Data pre-processing and analysis

#### Imaging data pre-processing

Videos were motion-corrected using a whole-frame cross-correlation algorithm algorithm^77^. Source extraction was done using the CaImAn package, which employs a constrained non-negative matrix factorization (CNMF) algorithm to identify putative neurons (ROIs)^78^. Because GCaMP7f was expressed under a pan-neuronal promoter in D1-tdTomato mice, tdTomato-negative cellular ROIs were classified as putative iSPNs rather than definitively identified iSPNs. We manually inspected all ROIs and excluded obvious non-cellular ROIs, neuropil-like ROIs, ROIs with ambiguous tdTomato overlap, and rare ROIs with morphology or fluorescence dynamics inconsistent with SPNs (Extended Data Fig. 1)^12^. Neurons with very high baseline fluorescence relative to other surrounding neurons, indicative of interneurons with high baseline firing rates, were also excluded (Extended Data Fig. 1D). We did not apply a quantitative interneuron classifier, and therefore cannot exclude rare interneuron contamination among tdTomato-negative ROIs. However, interneurons comprise only a small minority of rodent striatal neurons, and prior imaging studies indicate that interneuron calcium signals were typically lower amplitude and less likely to be detected by our ROI extraction and transient-selection procedures than SPN signals^12,37^. Most importantly, the main trajectory-dependent imbalance pattern was reproduced in an independent A2A-Cre cohort in which iSPNs were positively identified by tdTomato expression (Extended Data Fig. 3). Thus, rare interneuron contamination is unlikely to account for the population-level, trajectory-dependent dSPN/iSPN imbalance patterns reported here. In Drd1a-tdTomato mice, D1-positive neurons were identified by manually inspecting the alignment between the red tdTomato channel and ROI masks from the green GCaMP7f channel. Green ROI masks that clearly overlapped with a tdTomato-positive soma were classified as dSPNs, whereas ROIs that unambiguously did not overlap with a tdTomato-positive soma were classified as putative iSPNs (Extended Data Fig. 1A). In A2A-Cre mice, green ROI masks that clearly aligned with a tdTomato-positive soma were classified as putative iSPNs, whereas ROIs lacking red-channel overlap were classified as dSPNs (Extended Data Fig. 1B). Conversion to ΔF/F was done through the method provided by CalmAn, where the baseline fluorescence (F) was determined as the 8th quantile over a 500 frame moving window. ΔF/F was then thresholded to isolate significant positively going Ca^2+^ transients, defined as events exceeding 2 standard deviations above the median of a null distribution. The null distribution was made by truncating the session ΔF/F with normal distribution parameters in an exponentially modified Gaussian distribution, with parameters estimated using a maximum likelihood estimation (MLE) approach. Only sessions with more than 10 trials (VR-on tasks) or bouts (VR-off task) and more than 5 active dSPNs and iSPNs were included in all analyses (active cells were defined as more than 10 Ca^2+^ transients in the spontaneous running sessions or during track transversal in the virtual track tasks).

#### Behavioral variables and binning

The voltage output from the optical sensor was converted to linear velocity in m/s. Velocity traces were smoothed twice consecutively using the MATLAB ‘smooth’ function with window sizes of 1s and 0.5s. Acceleration was the derivative of the smoothed velocity, smoothed again using moving average with a 0.1 second window. Locomotion bouts were defined as periods in which velocity remained above 3 cm/s for more than 3.5 s and peak velocity exceeded 10 cm/s. Movement onset was defined as the first time point above the velocity threshold, and movement offset as the last time point above threshold. Total bout distance was calculated as the cumulative distance traveled from 1 s before movement onset to 1 s after movement offset. For VR-on tasks, only bout onsets and offsets occurring outside the inter-trial interval were included in event-triggered analyses. For spontaneous running outside of VR, bouts were excluded if peak velocity did not exceed 15 cm/s, to enable comparison with the in-VR velocity range, or if reward delivery occurred at any point during bout progression. For the infinite-track task, analyses were restricted to bouts longer than 150 cm with peak velocity exceeding 15 cm/s.

For position-based analyses in VR-on tasks, the main track, defined from the end of the start zone to the reward location, was divided into 100 position bins. For the VR-off task, position was normalized to the percentage of total distance traveled within each bout. For the infinite-track task, the repeating virtual track was divided into 100 equal position bins for each trial, while for distance-based analyses, the first 150 cm of each selected bout was divided into 100 equal distance bins. To account for variability between mice and session and repeated measures, we fit the position binned or triggered average velocity or acceleration to a Generalized Linear Mixed Effect model (GLME) (see below) using the MATLAB function iitglme^1^. The model considers mice and sessions as random effects, with sessions nested within mice. The estimated population mean and the confidence interval were then used to plot the center line and shaded regions of the line plots.

#### Quantification of mean population activity and dSPN/iSPN differences

For plots showing mean ΔF/F or differences between dSPN/iSPN ΔF/F means, we first calculated the means for each trial (binned by position or triggered on events as indicated in each figure) and across each subpopulation (as indicated in each figure, dSPNs/iSPNs, tuned/untuned, etc.) for each imaging field. In addition to showing mean ΔF/F transients, we also characterized mean population activity using: 1) the percentage of neurons active over time, calculated at each time point as the fraction of putative iSPNs or dSPNs whose unthresholded (non-transient) ΔF/F exceeded 2 standard deviations above that neuron’s mean activity across the session; and 2) estimated event rates deconvolved from the non-transient ΔF/F traces (Extended Data Fig. 4A-H), obtained using a non-negative FOOPSI deconvolution algorithm with a first-order autoregressive model (AR(1), **p**=1). Deconvolution was implemented in the unconstrained formulation, with the calcium decay time constant fixed at 0.7 s based on prior recommendations for the fast GCaMP7f sensor^79,80^. To calculate the mean activity of dSPNs and iSPNs or their difference, accounting for variability across trials, sessions, and mice, we used a Generalized Linear Mixed Effect model (MATLAB function ‘fitglme’) for each bin or timepoint with the following equation:

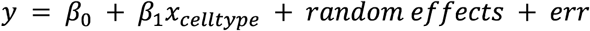

For a given binary condition variable *x*, *β*_0_ is the estimated mean given *x* = 0, and *β*_1_ is the estimated difference between the two level of *x* (dSPN and iSPN). Unless otherwise specified, the mixed effect model considers mice and sessions as random effects, with sessions nested within mice. The 95% confidence intervals of the mean or difference terms were displayed as shaded regions in all plots.

#### Position tuning and track sensitivity

To determine whether individual neurons were stably tuned to a task-specific representation within a session, we compared mean binned ΔF/F maps computed from odd and even trials. For position-tuning analyses in the finite and infinite track tasks, ΔF/F was binned by track position to generate position rate maps. For distance-tuning analyses in the infinite track task, ΔF/F was binned by distance traveled within each locomotion bout at a fixed bin size of 1.5 cm to generate distance rate maps. The odd-even correlation for each neuron was then compared to a null distribution generated by randomly circularly shifting that neuron’s ΔF/F relative to trial epochs by more than 33 ms over 500 iterations, followed by rebinning and recomputing the odd-even correlation. Neurons were classified as significantly tuned (to either location or distance) if the true correlation exceeded the 95th percentile of the shuffled distribution and the neuron exhibited significant ΔF/F transients on at least 40% of trials or bouts. For significantly tuned neurons, field width was defined by identifying the bin with the maximum ΔF/F and then finding the bins on either side where ΔF/F fell below 20% of the difference between the maximum and minimum ÄF/F values.

Track sensitivity was assessed in sessions with two tracks. For each neuron, mean position rate maps were computed separately for track 1 and track 2, and the true cross-track correlation was compared to a bootstrap distribution generated by randomly swapping trials between tracks over 500 iterations. Neurons were classified as track position sensitive if the true cross-track correlation was lower than the 1st percentile of the randomized distribution.

To compare track-sensitive, track-insensitive, and untuned population in the two-track task (Extended Data Fig. 7K-M), track-sensitive and track-insensitive populations were restricted to neurons with significant position tuning on the most familiar track (i.e., the track experienced before introduction of the second environment). Only trials from the most familiar track were included in these plots to avoid confounds from learning– or novelty-dependent effects in the less familiar track. Untuned neurons were included only if ΔF/F exceeded 0.5 on at least 40% of trials, to control for overall activity differences introduced by the tuning criteria. For infinite-track subpopulation analyses, the distance-tuned population included neurons with significant distance tuning but no significant position tuning, whereas the non-distance-tuned population included neurons tuned only to position or untuned to either representation, provided that ΔF/F exceeded 0.5 on at least 40% of locomotion bouts.

#### Computational model

We developed a simplified computational model to test whether structured differences between direct– and indirect-pathway striatal populations (dSPNs and iSPNs) could emerge through learning on position-related representations and, in a feedback extension, reshape locomotor trajectories. The model was designed to capture three features of the data: mixed position-related and locomotor inputs in individual neurons, opponent plasticity in dSPN and iSPN populations, and a kinematics-linked teaching signal derived from locomotor acceleration.

The network contained 1,000 model neurons (500 dSPNs and 500 iSPNs). Cell identity was represented by a pathway sign variable, *s_i_* = +1for dSPNs and *s_i_* = —1for iSPNs. Simulations were run in discrete time (Δ *t* = 0.05 s) on a 1.5 m track, and position-dependent quantities were evaluated in 2-cm bins. Each neuron received a pseudorandom mixture of position-related and locomotor inputs whose weighted sum determined its activity. Position-related input was represented by a localized Gaussian field tiling the trajectory, whereas locomotor input included velocity and positive acceleration terms. Initial synaptic weights were drawn from the same distributions for dSPN and iSPN populations, such that pathway activity was approximately balanced before learning and no structured dSPN/iSPN imbalance was imposed a priori.

At each time step, neural activity was computed as a rectified sum of position-related drive, locomotor drive, baseline activity, and Gaussian output noise:

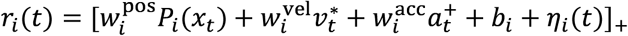

where *P_i_*(*x_t_*)is the neuron’s position field, 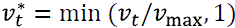 is normalized velocity, 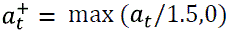is normalized positive acceleration, *b_i_* is baseline activity, and *n_i_*(t)is additive Gaussian noise. Default parameters were *n*_trials_ = 150, σ_pos_ = 0.12m, *v*_max_ = 0.35m/s, baseline mean = 0.02, baseline s.d. = 0.01, and output-noise s.d. = 0.02.

Position-related synapses were updated using an eligibility-trace rule driven by a temporally filtered, delayed signed-acceleration signal. The eligibility trace evolved as

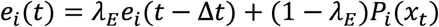

where 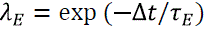 and 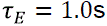. The teaching signal was derived from normalized signed acceleration, 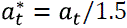, which was first low-pass filtered,

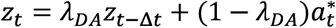

with 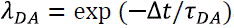 and 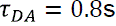, and then delayed by 200 ms:

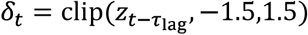

with 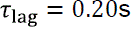. Position weights were updated according to

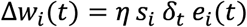

and clipped to the interval [0, 1]. This imposed lag is consistent with measured acceleration-related dopamine signals in previous studies. Thus, positive teaching signals strengthened dSPN-associated synapses and weakened iSPN-associated synapses, whereas negative signals produced the opposite update. In all manuscript figures, plasticity was restricted to position-related inputs, whereas generic locomotor inputs remained fixed (see Discussion).

#### Learning model for emergence of imbalance

To test how structured imbalance could emerge through learning, locomotor kinematics were imposed externally. On each trial, motion began at the start of the track and followed a prescribed trajectory with acceleration, cruising, and deceleration phases before reward. Trial-to-trial variability was introduced by sampling maximum velocity and the lengths of the acceleration and deceleration phases from bounded Gaussian distributions. Values were *v*_max_ = 0.20 土 0.02m/s, acceleration length = 0.20 ± 0.03m, and deceleration length = 0.20 ± 0.03m. Reward was delivered at the end of the track, followed by a 3-s post-reward dwell. This model was used to test whether opponent plasticity acting on initially balanced populations could generate structured dSPN/iSPN imbalance across the trajectory.

#### Feedback model for restructuring locomotion

To test whether learned imbalances could also influence behavior, we implemented a feedback extension in which SPN imbalance biased a locomotion controller (Fig. 6H). In this model, behavior was initially governed by an exploratory controller that generated variable trajectories with acceleration at bout onset and stopping near the reward zone. Neural activity was computed as above, using one position field per neuron and plasticity on position weights only.

At each time step, a local imbalance signal,

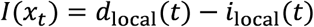

was computed from the difference between local dSPN and iSPN population activity determined by the currently active position fields. Rather than acting instantaneously on velocity, this signal was transformed into a locomotion controller bias in two stages. First, imbalance was low-pass filtered into a smoothed drive state,

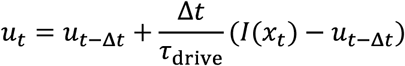

and then into a more persistent bias state,

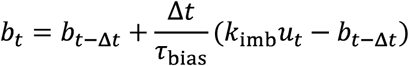

with clipping to a bounded range. Here, *k*_imb_ is the gain from SPN imbalance to controller bias, *u_t_* is the smoothed imbalance drive, and *b_t_* is the persistent bias term. Target velocity was then given by

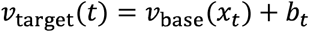

The baseline controller included a constant cruising term together with an additional slowing term in the reward zone. Actual velocity relaxed toward the target with first-order dynamics and additive Gaussian noise:

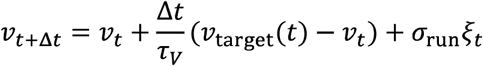

Before reward, velocity was constrained to remain above a small positive minimum to prevent premature stopping; after reward, it decayed to zero during the dwell period. Default closed-loop parameters were *v*_CruiseBase_ = 0.16m/s, 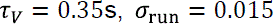, reward-zone onset = 1.40m, reward-zone slowing gain =0.08, *k*_imb_ =0.13, 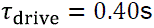, and 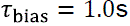. The same acceleration-derived teaching signal used in the learning model updated position weights in the feedback model.

#### Analysis of simulated activity

Simulated neural activity, velocity, and acceleration were analyzed by binning values as a function of track position. In the learning model, representative heat maps were generated from single simulations, whereas average curves were summarized across 100 independent simulations. In the feedback model, representative trial-by-trial velocity plots were generated from one simulation, and summary metrics were computed from the first and last 10 trials of each simulation and summarized across 25 independent simulations. Trial-to-trial stereotypy was quantified as the mean Pearson’s correlation between individual velocity profiles and the simulation-specific template trajectory. Sustained early velocity was quantified as normalized velocity AUC over the first half of the track. Braking position was defined as the position at which velocity entered a sustained late decline, and slowing magnitude was defined as the drop in velocity between a late plateau window and the end of the track.

For display of representative neuronal heat maps, the strongest position-tuned neurons were identified from the post-learning modulation depth of their position-binned activity curves, normalized on a per-neuron basis to the interval [0, 1], and sorted by the position bin of peak activity. This normalization and sorting were used for visualization only and did not affect summary analyses.

Notes on *statistical tests and sample* sizes. To test whether the mean ΔF/F difference between dSPNs and iSPNs differed significantly across experimental conditions (e.g. Extended Data Fig. 5K-M), an interaction term between cell type and experimental condition was added to the previous GLME model:

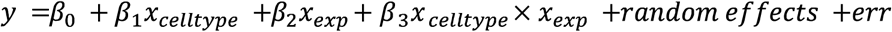

where *β_3_* represents the change in the mean dSPN-iSPN difference between conditions. Non-parametric tests for significance were used for all analyses unless otherwise noted. Specific tests and sample sizes are indicated within figure legends or in the main text. Multiple comparisons were corrected using the Bonferroni-Holm’s correction. (Bonferroni-Holm Correction for Multiple Comparisons – File Exchange – MATLAB Central (mathworks.com))

## Acknowledgements

This work was supported by a Klingenstein-Simons’s Foundation fellowship, Whitehall Foundation Fellowship, National Institute of Mental Health (R01 MH125835) to M.W.H and a Postdoctoral Training Fellowship to B.F. through the Boston University Center for Systems Neuroscience. We thank the Boston University Centers for Neurophotonics and Systems Neuroscience for financial and technical support and Boston University Animal Science Center for providing central laboratory and animal care and support resources. We thank Dr. Mai-Anh Vu, Ryan Senne and Kylie Feliciano for their technical help with analysis pipelines, task design, and equipment setup. We thank Drs. Milan Valyear and Eleanor Brown for their comments on a draft of the manuscript. The mouse A2A-cre mouse strain (B6.FVB(Cg)-Tg(Adora2a-cre)KG139Gsat/Mmucd, RRID:MMRRC_036158-UCD), was obtained from the Mutant Mouse Resource and Research Center (MMRRC) at University of California at Davis, an NIH-funded strain repository, and was donated to the MMRRC by MMRRC at University of California, Davis. Made from the original strain (MMRRC:034637) donated by Nathaniel Heintz, Ph.D., The Rockefeller University, GENSAT and Charles Gerfen, Ph.D., National Institutes of Health, National Institute of Mental Health

---

**Extended Data Fig. 1:**
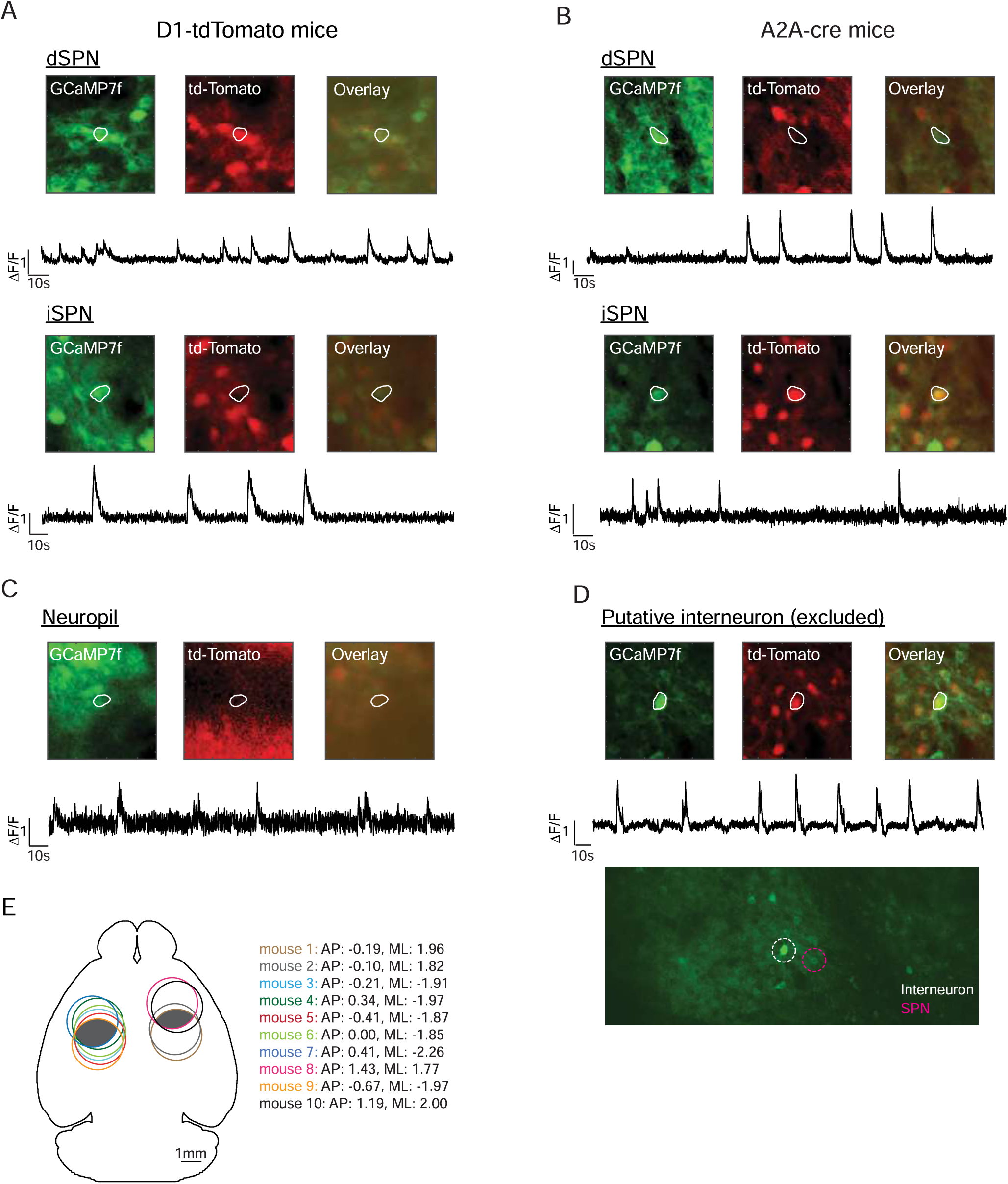
Cell-type identification, ROI exclusion criteria, and cannula placement for dorsal striatal imaging. **A**, Example ROI classification in D1-tdTomato mice. GCaMP7f fluorescence, tdTomato fluorescence, and merged images are shown for an example tdTomato-positive cellular ROI classified as a dSPN and an example tdTomato-negative cellular ROI classified as a putative iSPN. White outlines indicate the selected ROI, and traces show example ΔF/F activity from the outlined ROI. B, Example ROI classification in A2A-Cre mice with tdTomato expression in iSPNs. GCaMP7f fluorescence, tdTomato fluorescence, and merged images are shown for an example tdTomato-negative cellular ROI classified as a putative dSPN and an example tdTomato-positive cellular ROI classified as an iSPN. White outlines indicate the selected ROI, and traces show example ΔF/F activity from the outlined ROI. C, Example ROI classified as neuropil and excluded from analysis because it lacked a clearly delineated cellular morphology. White outline indicates the excluded ROI, and the trace shows example ΔF/F from that ROI. D, Example ROI excluded from analysis due to its bright baseline F relative to surrounding neurons. The morphology and activity pattern are consistent with striatal GABAergic interneurons. E, Approximate cannula positions estimated from post-hoc histological inspection and projected onto a horizontal brain outline. Only a subset of imaged mice with histology available are shown. Colored outlines indicate estimated cannula positions for individual mice; darker shaded region indicates the intersection of all placements. Coordinates indicate estimated cannula position centers relative to bregma.

**Extended Data Fig. 2:**
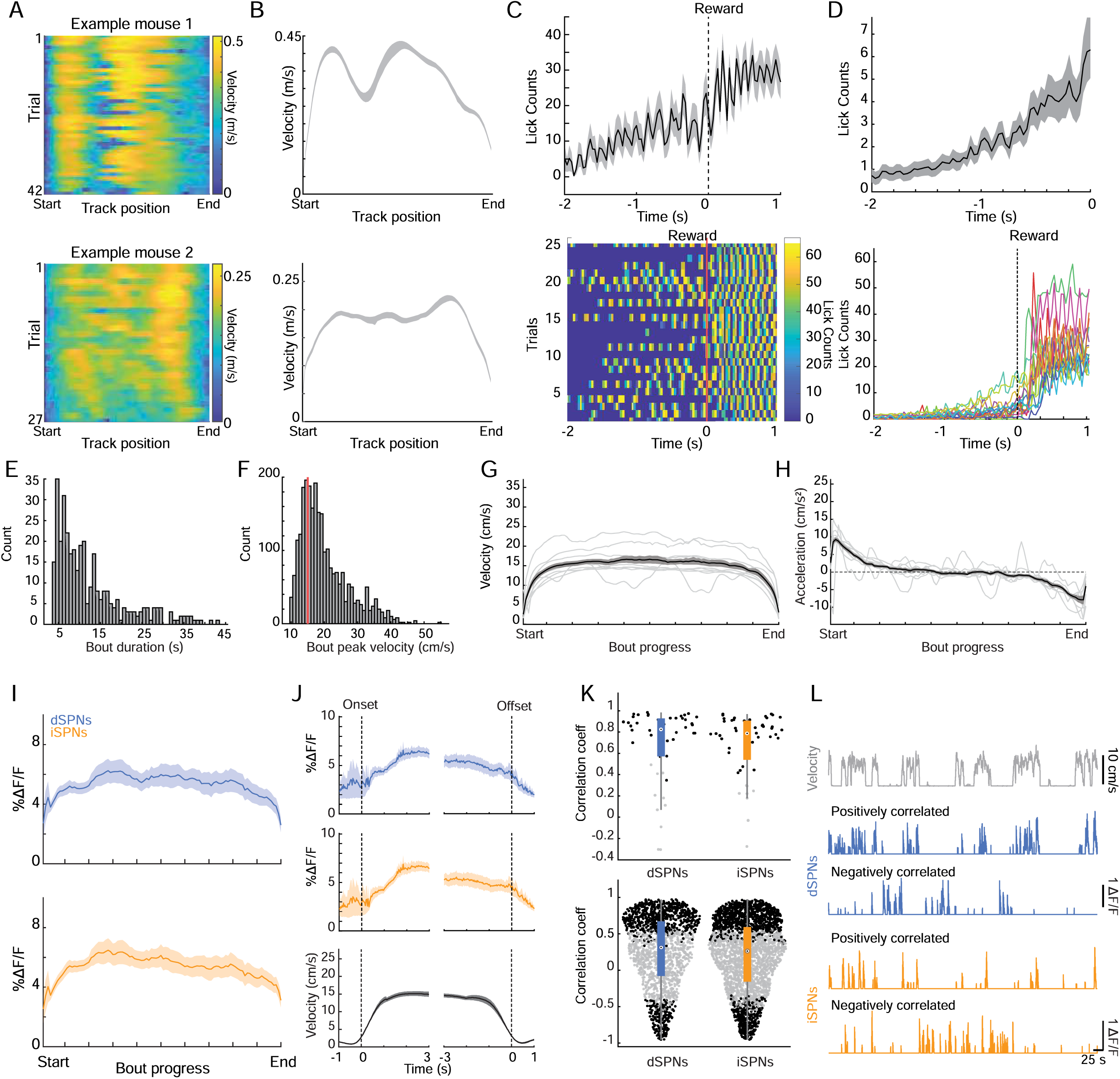
Licking, behavioral bout selection, and spontaneous-locomotion activity outside the virtual track task. **A**, Velocity rasters from locomotor bouts during virtual track traversal for single representative sessions in two mice. Each row is velocity binned by track position for one trial. B_r_ Mean velocity profiles for the example sessions shown in A. C, Top, mean lick count aligned to reward delivery during the virtual track task from one representative session. Bottom, trial-by-trial lick counts for the same session. D, Licking aligned to reward delivery during the virtual track task. Top, model-estimated mean lick counts across mice and sessions. Bottom, individual mouse averages. E, Distribution of spontaneous locomotor bout durations outside VR. F, Distribution of spontaneous locomotor bout peak velocities outside VR. Red line indicates the peak bout-velocity cutoff of 15 cm/s used to match spontaneous locomotor bouts to track-traversal bouts. G, Mean velocity binned by normalized bout progress during spontaneous locomotion outside VR (n = 9 mice, 41 sessions, 345 bouts). Gray traces show individual sessions and black trace indicates the model-estimated mean. H, Mean acceleration binned by normalized bout progress during spontaneous locomotion outside VR (n = 9 mice, 41 sessions, 345 bouts). Gray traces show individual sessions and black trace indicates the model-estimated mean. I, Mean population ΔF/F of dSPNs and iSPNs binned by normalized bout progress during spontaneous locomotion outside VR (n = 9 mice, 41 sessions, 48 imaging fields; 1914 dSPNs, 3248 iSPNs). **J_f_** Mean dSPN activity, iSPN activity, and velocity aligned to locomotor onset and offset during spontaneous locomotion outside VR. Dashed vertical lines indicate onset or offset times. **K,** Correlations between dSPN or iSPN activity and locomotor velocity during spontaneous locomotion outside VR. Top, correlations between population activity binned by velocity at a fixed bin size of 0.05 cm/s and binned velocity across imaging fields (n = 9 mice, 41 sessions, 48 imaging fields; significant fields: dSPN, 79.2%; iSPN, 83.3%; positive among significant fields: dSPN, 100%; iSPN, 100%). Bottom, velocity-binned single-neuron activity correlations with velocity (n = 1914 dSPNs, 3248 iSPNs; significant neurons: dSPN, 45.2%; iSPN, 39.7%; positive among significant neurons: dSPN, 77.6%; iSPN, 72.8%). Black points indicate significant correlations and gray points indicate non-significant correlations. Box plots show the median and interquartile range of correlation coefficients; individual points indicate imaging fields or neurons as indicated. L, Example dSPNs and iSPNs with positive or negative correlations with locomotor velocity during spontaneous locomotion outside VR. Shaded regions in all line plots are the 95% confidence intervals of the model coefficients from the linear mixed-effects model.

**Extended Data Fig. 3:**
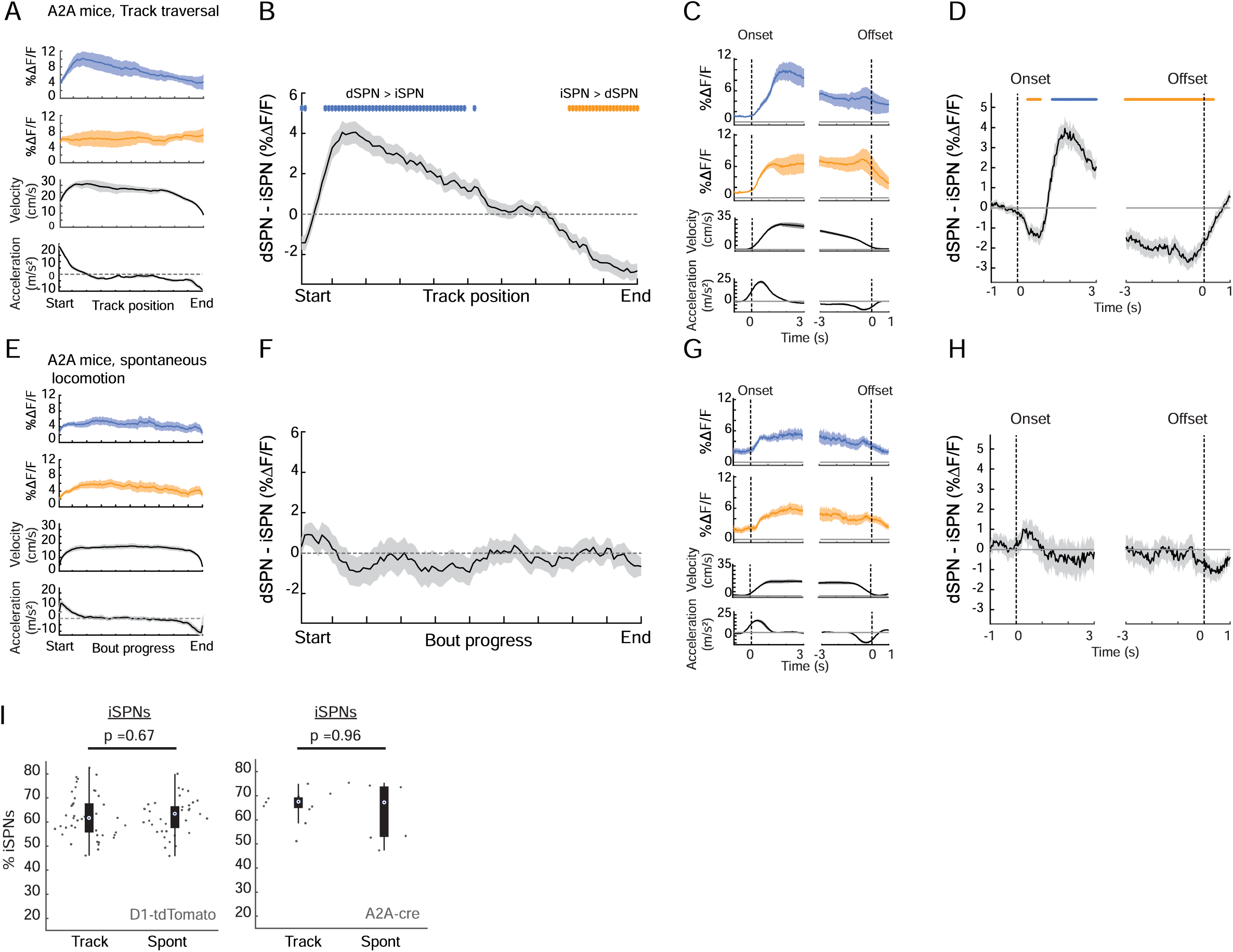
Track-related dSPN/iSPN imbalances are robust to cell-type labeling strategy. **A**, Mean ΔF/F of dSPNs, iSPNs, velocity, and acceleration binned by track position during track traversal in A2A-Cre mice with tdTomato expression in iSPNs (n = 4 mice, 12 sessions/imaging fields; 159 dSPNs, 310 iSPNs). B_r_ Mean dSPN-iSPN activity difference binned by track position during track traversal in A2A-Cre mice. Positive values indicate greater dSPN activity and negative values indicate greater iSPN activity. C, Mean dSPN activity, iSPN activity, velocity, and acceleration aligned to locomotor onset and offset during track traversal in A2A-Cre mice. Dashed vertical lines indicate onset or offset times. D, Mean dSPN-iSPN activity difference aligned to locomotor onset and offset during track traversal in A2A-Cre mice. **E,** Mean ΔF/F of dSPNs, iSPNs, velocity, and acceleration binned by normalized bout progress during spontaneous locomotion outside the virtual track task in A2A-Cre mice (n = 3 mice, 8 sessions/imaging fields; 132 dSPNs, 236 iSPNs). F, Mean dSPN-iSPN activity difference binned by normalized bout progress during spontaneous locomotion in A2A-Cre mice. G, Mean dSPN activity, iSPN activity, velocity, and acceleration aligned to locomotor onset and offset during spontaneous locomotion in A2A-Cre mice. H, Mean dSPN-iSPN activity difference aligned to locomotor onset and offset during spontaneous locomotion in A2A-Cre mice. I, Fraction of cells classified as iSPNs in D1-tdTomato mice and A2A-Cre mice during track traversal and spontaneous locomotion. Each point indicates one imaging field. Box plots show the median and interquartile range of cell-type fractions. P values indicate comparisons between track traversal and spontaneous locomotion within each cohort (Wilcoxon rank sum test). Blue and orange bars indicate position bins or time points with significant dSPN > iSPN or iSPN > dSPN differences, respectively. Shaded regions in all line plots are the 95% confidence intervals of the model coefficients from the linear mixed-effects model. Significance bars indicate p < 0.05, t-tests on model coefficients, Bonferroni corrected for multiple comparisons.

**Extended Data Fig. 4:**
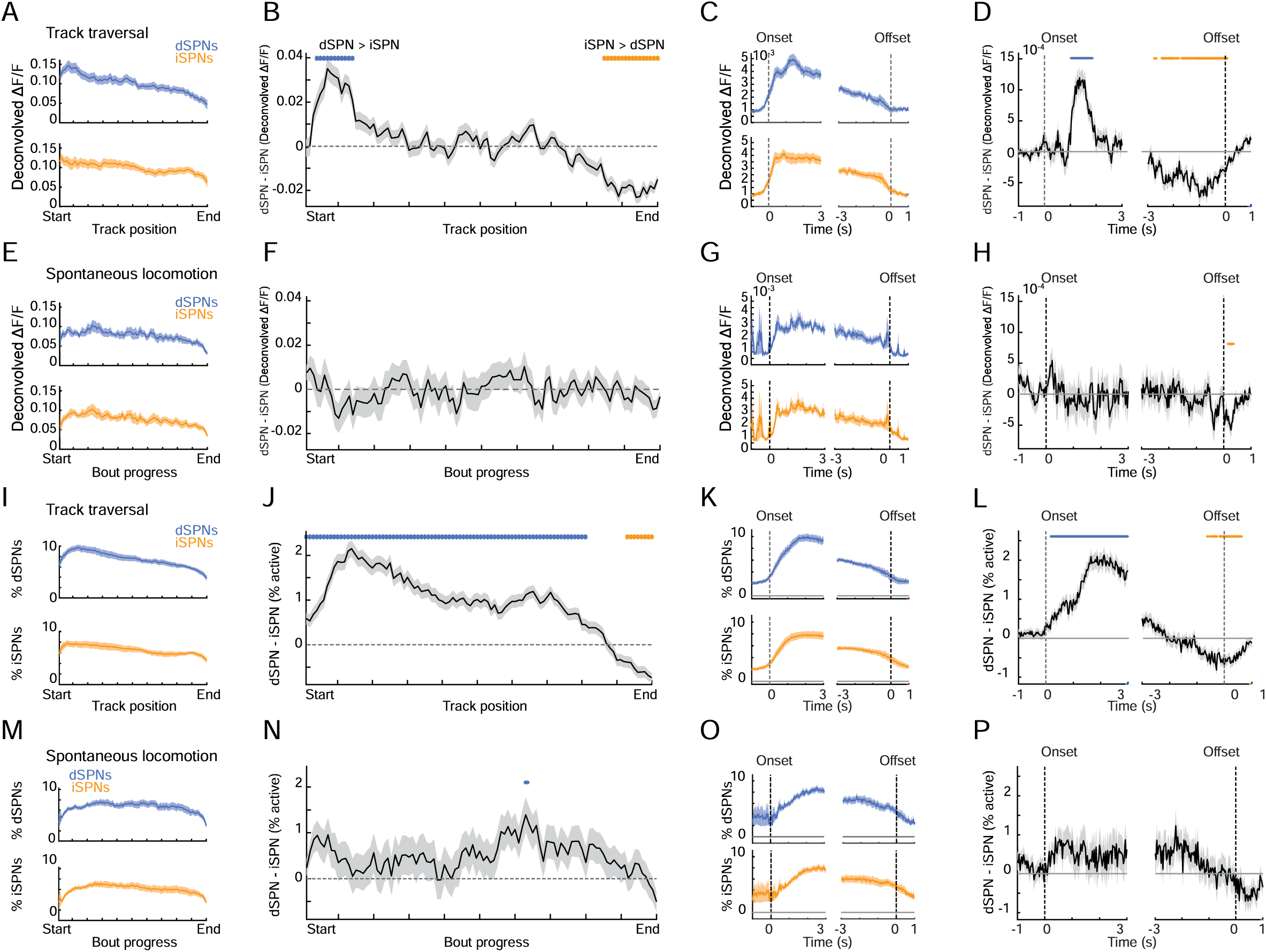
Track-related dSPN/iSPN imbalances are reproduced using deconvolved activity and active-cell fractions. **A-D**, Deconvolved activity analyses during track traversal (n = 15 mice, 55 sessions; 1770 dSPNs, 2809 iSPNs). A, Mean deconvolved activity of dSPNs and iSPNs binned by track position during track traversal. B, Mean dSPN-iSPN difference in deconvolved activity binned by track position during track traversal. C, Mean deconvolved dSPN and iSPN activity aligned to locomotor onset and offset during track traversal. D, Mean dSPN-iSPN difference in deconvolved activity aligned to locomotor onset and offset during track traversal. **E-H,** Deconvolved activity analyses during spontaneous locomotion outside the track task (n = 9 mice, 41 sessions, 48 imaging fields; 1914 dSPNs, 3248 iSPNs). E, Mean deconvolved activity of dSPNs and iSPNs binned by normalized bout progress during spontaneous locomotion. **F,** Mean dSPN-iSPN difference in deconvolved activity binned by normalized bout progress during spontaneous locomotion. G, Mean deconvolved dSPN and iSPN activity aligned to locomotor onset and offset during spontaneous locomotion. H, Mean dSPN-iSPN difference in deconvolved activity aligned to locomotor onset and offset during spontaneous locomotion. I-P, Active-cell fraction analyses from the same mice, sessions, imaging fields, and neurons shown in A-H. I, Fraction of active dSPNs and iSPNs binned by track position during track traversal. J_f_ Difference between the fraction of active dSPNs and iSPNs binned by track position during track traversal. **K,** Fraction of active dSPNs and iSPNs aligned to locomotor onset and offset during track traversal. L, Difference between the fraction of active dSPNs and iSPNs aligned to locomotor onset and offset during track traversal. M_r_ Fraction of active dSPNs and iSPNs binned by normalized bout progress during spontaneous locomotion. N, Difference between the fraction of active dSPNs and iSPNs binned by normalized bout progress during spontaneous locomotion. **O,** Fraction of active dSPNs and iSPNs aligned to locomotor onset and offset during spontaneous locomotion. P, Difference between the fraction of active dSPNs and iSPNs aligned to locomotor onset and offset during spontaneous locomotion. Blue and orange bars indicate position bins or time points with significant dSPN > iSPN or iSPN > dSPN differences, respectively. Shaded regions in all line plots are the 95% confidence intervals of the model coefficients from the linear mixed-effects model. Box plots show the median and interquartile range. Significance bars in A-P indicate p < 0.05, t-tests on model coefficients, Bonferroni corrected for multiple comparisons.

**Extended Data Fig. 5:**
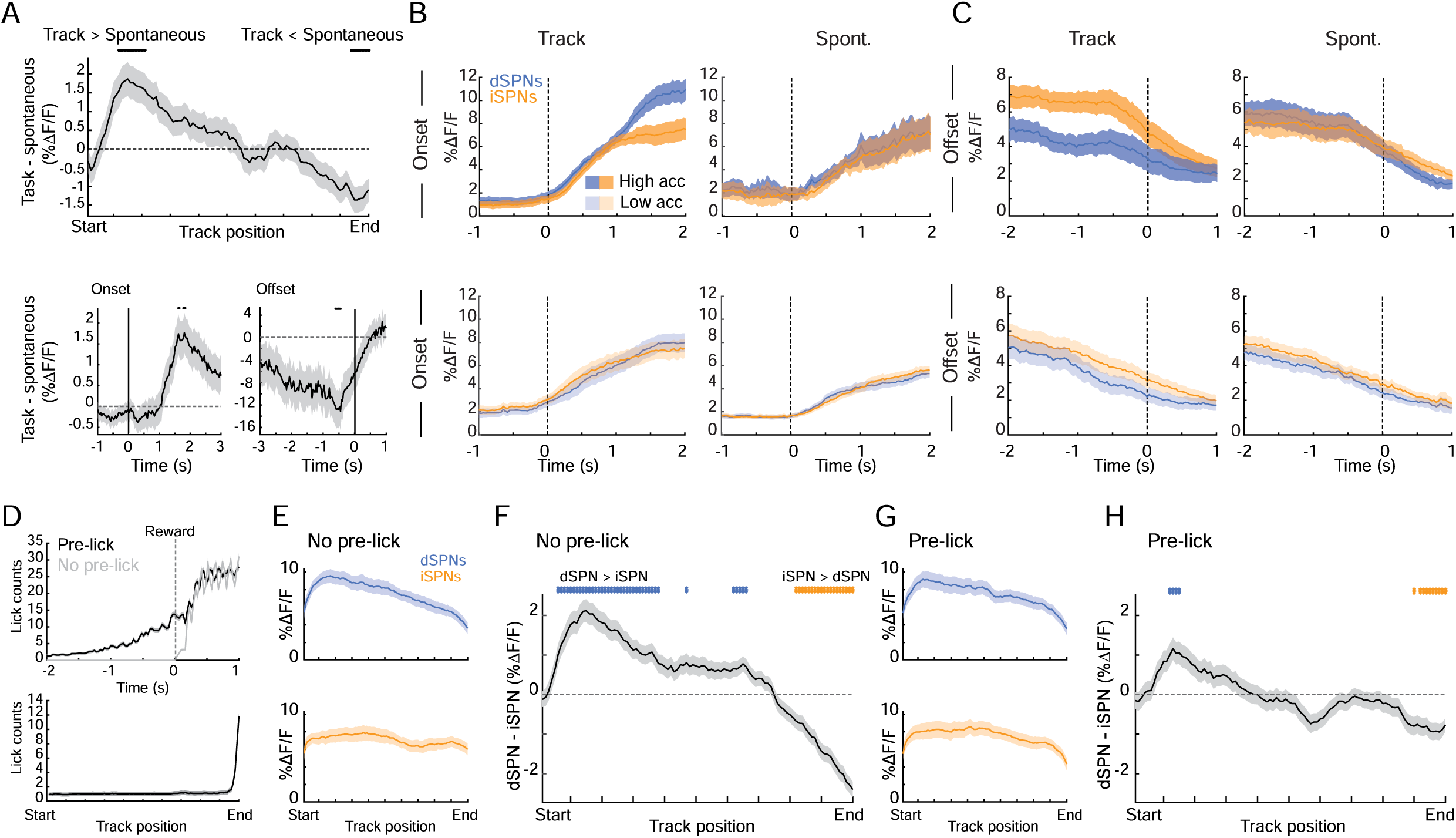
dSPN/iSPN imbalances are not present during spontaneous locomotion or dependent upon anticipatory licking in the task. **A**, Difference between dSPN/iSPN imbalances during track traversal and spontaneous locomotion, corresponding to the comparisons shown in Fig. 2A-D. Top, track-position-binned difference between track traversal and spontaneous locomotion. Bottom, onset– and offset-aligned differences between track traversal and spontaneous locomotion. Positive values indicate a larger dSPN-favored imbalance during track traversal than during spontaneous locomotion, and negative values indicate a larger iSPN-favored imbalance during track traversal than during spontaneous locomotion (n = 16 mice, 96 sessions, 106 imaging fields). B, Mean dSPN and iSPN activity aligned to locomotor onset for high– and low-acceleration groups during track traversal and spontaneous locomotion, corresponding to the imbalance comparisons shown in Fig. 2F,G. Top row, high-acceleration groups; bottom row, low-acceleration groups. C, Mean dSPN and iSPN activity aligned to locomotor offset for high– and low-deceleration groups during track traversal and spontaneous locomotion, corresponding to the imbalance comparisons shown in Fig. 2I,J. Top row, high-deceleration groups; bottom row, low-deceleration groups. D, Licking behavior on trials with and without anticipatory licking before reward delivery during track traversal. Top, lick counts aligned to reward delivery. Bottom, lick counts binned by track position for trials with anticipatory licking and trials without anticipatory licking (with anticipatory licking: n = 15 mice, 53 sessions, 711 trials; without anticipatory licking: n = 15 mice, 55 sessions, 763 trials). E, Mean dSPN and iSPN activity binned by track position on trials without anticipatory licking. F, Mean dSPN-iSPN activity difference on trials without anticipatory licking. G, Mean dSPN and iSPN activity binned by track position on trials with anticipatory licking. H, Mean dSPN-iSPN activity difference on trials with anticipatory licking. Blue and orange bars indicate position bins or time points with significant dSPN > iSPN or iSPN > dSPN differences, respectively. Black bars indicate time points or position bins with significant differences between track traversal and spontaneous locomotion in A. Shaded regions in all line plots are the 95% confidence intervals of the model coefficients from the linear mixed-effects model. Significance bars indicate p < 0.05, t-tests on model coefficients, Bonferroni corrected for multiple comparisons.

**Extended Data Fig. 6:**
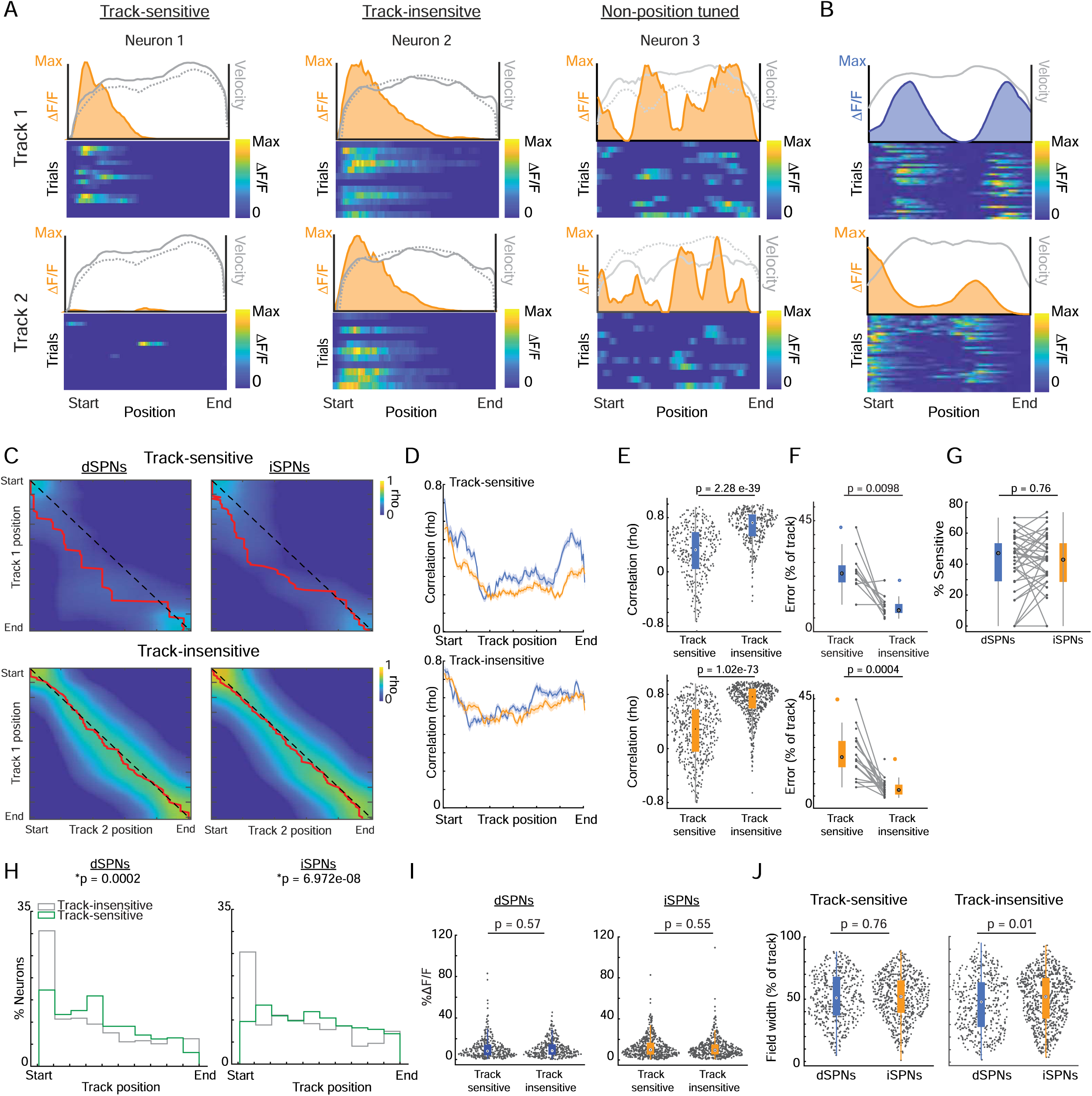
Characterization of track-sensitive and track-insensitive position tuning. **A**, Representative iSPNs classified as track-sensitive, track-insensitive, or non-position tuned. Line plots show mean ΔF/F in Track 1 and Track 2, with overlaid velocity traces in gray. Heatmaps show single-trial ΔF/F binned by track position. B_r_ Representative dSPN (top) and iSPN (bottom) examples classified as tuned and track-sensitive that exhibited multiple activity fields. Line plots and heatmaps are as in A. C, Pairwise correlations (Spearman’s rho) between mean ΔF/F population vectors at each Track 1 position and mean ΔF/F population vectors at each Track 2 position, computed across track-sensitive (top) or track-insensitive (bottom) dSPNs and iSPNs (track-sensitive neurons tuned to Track 1: n = 202 dSPNs, 280 iSPNs; Track 2: n = 251 dSPNs, 341 iSPNs; track-insensitive neurons: n = 327 dSPNs, 559 iSPNs). Matrices show correlations for all combinations of Track 1 and Track 2 positions. Values along the diagonal dashed line indicate correlations between the same relative positions in the two tracks, such that high diagonal values indicate similar population activity patterns at corresponding positions. Red lines indicate, for each Track 1 position, the Track 2 position with the highest correlation. D_r_ Mean correlations between Track 1 and Track 2 population vectors at corresponding relative positions, corresponding to the diagonal values in C. Correlations were computed for each session at each position and then averaged across sessions for track-sensitive (top) and track-insensitive (bottom) dSPNs and iSPNs. E, Correlations between position-binned mean ΔF/F in Track 1 and Track 2 for each neuron classified as track-sensitive or track-insensitive. Each point indicates one neuron; box plots show the median and interquartile range of correlation coefficients (track-sensitive: n = 332 dSPNs, 494 iSPNs; track-insensitive: n = 327 dSPNs, 559 iSPNs). P values indicate comparisons between track-sensitive and track-insensitive neurons within each cell type (Wilcoxon rank sum test). F, Absolute deviation, or spatial error, between the empirically observed peak correlation position and the expected peak correlation position if activity patterns were identical between Track 1 and Track 2, calculated from the distance between the red line and unity diagonal in C. Each point indicates one session with more than 10 dSPNs or iSPNs from either tuning class; paired lines connect track-sensitive and track-insensitive values from the same session (track-sensitive: n = 11 fields for dSPNs, 18 fields for iSPNs; track-insensitive: n = 13 fields for dSPNs, 20 fields for iSPNs). P values indicate comparisons between track-sensitive and track-insensitive populations within each cell type (paired, two-sided Wilcoxon signed-rank tests). G, Percentage of position-tuned dSPNs and iSPNs classified as track-sensitive. Each point indicates one session; paired lines connect dSPN and iSPN values from the same session (n = 39 fields). P value indicates comparison between dSPNs and iSPNs (paired, two-sided Wilcoxon signed-rank test). H, Distribution of field centers for track-sensitive and track-insensitive position-tuned dSPNs and iSPNs. P values indicate comparisons between field-center distributions for track-sensitive and track-insensitive neurons within each cell type (Kolmogorov-Smirnov test). **I,** Mean ΔF/F across the track for track-sensitive and track-insensitive dSPNs and iSPNs. Each point indicates one neuron; box plots show the median and interquartile range of mean ΔF/F. P values indicate comparisons between track-sensitive and track-insensitive neurons within each cell type (Wilcoxon rank sum test). J, Position-field width of dSPNs and iSPNs classified as track-sensitive or track-insensitive. Each point indicates one neuron; box plots show the median and interquartile range of field width. P values indicate dSPN versus iSPN comparisons within each tuning class (Wilcoxon rank sum test). Shaded regions in D are the standard error of session averaged correlation at each spatial location. Significance values are shown above each comparison.

**Extended Data Fig. 7:**
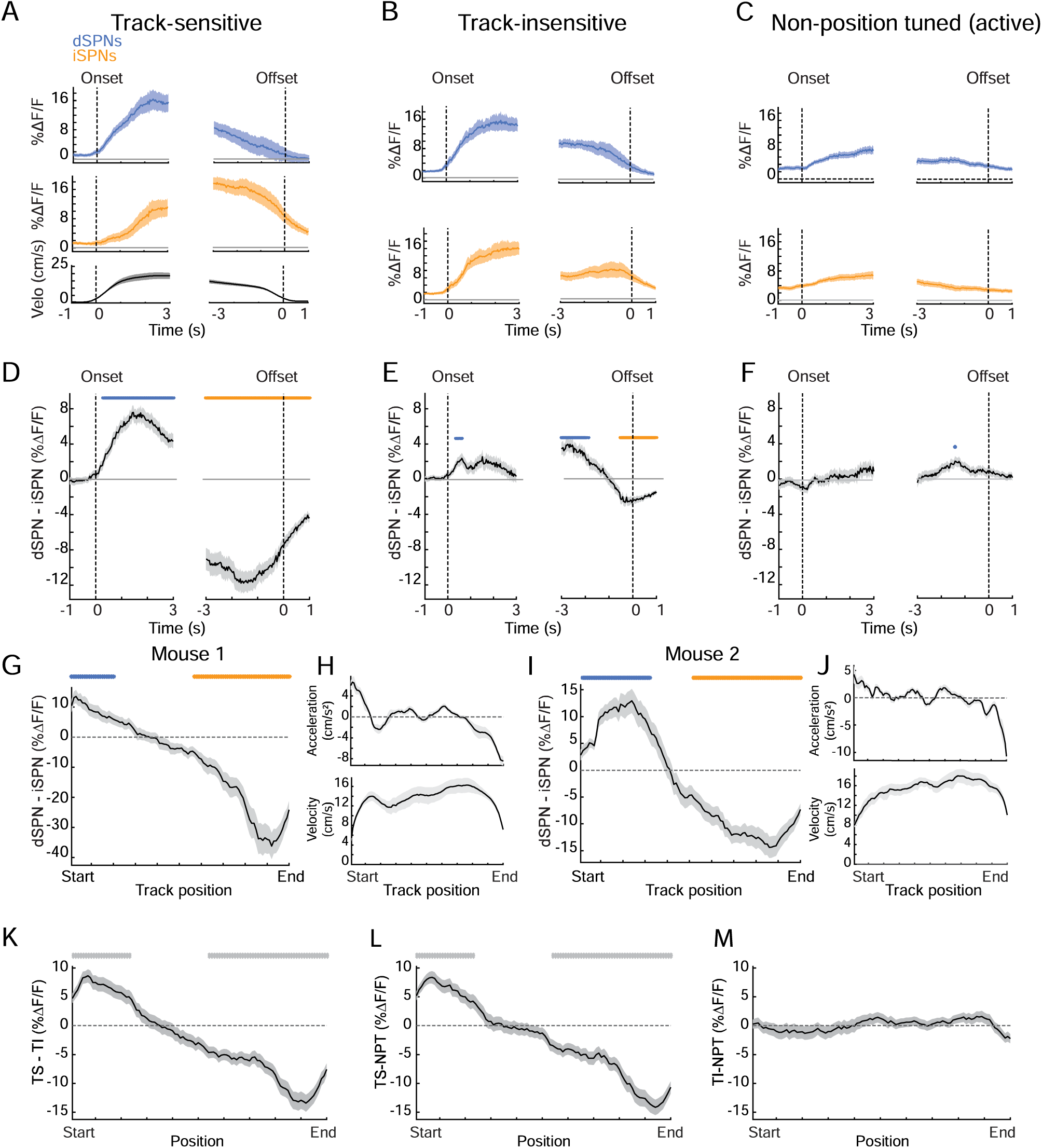
Additional aligned analyses and population contrasts for track-sensitive, track-insensitive, and non-position-tuned SPN populations. **A**, Mean dSPN activity, iSPN activity, and velocity aligned to locomotor onset and offset for track-sensitive, position-tuned neurons (n = 12 mice, 39 sessions; 202 dSPNs, 280 iSPNs). B_r_ Same as A for track-insensitive, position-tuned neurons (n = 12 mice, 39 sessions; 327 dSPNs, 559 iSPNs). **C,** Same as A for active non-position-tuned neurons (n = 12 mice, 34 sessions, 39 imaging fields; 251 dSPNs, 587 iSPNs). D, Mean dSPN-iSPN activity difference aligned to locomotor onset and offset for the track-sensitive population shown in A. E, Same as D for the track-insensitive population shown in B. F, Same as D for the active non-position-tuned population shown in C. **G,** Example dSPN-iSPN imbalance across track position from one mouse with a large number of track-sensitive cells. H, Acceleration and velocity profiles for the mouse shown in G. I, Example dSPN-iSPN imbalance across track position from a second mouse with a large number of track-sensitive cells. **J,** Acceleration and velocity profiles for the mouse shown in I. **K,** Difference between the mean dSPN-iSPN imbalance traces of the track-sensitive and track-insensitive populations across track position (TS-TI). L, Difference between the mean dSPN-iSPN imbalance traces of the track-sensitive and non-position-tuned populations across track position (TS-NPT). **M_r_** Difference between the mean dSPN-iSPN imbalance traces of the track-insensitive and non-position-tuned populations across track position (TI-NPT). TS, track-sensitive position-tuned population; Tl, track-insensitive position-tuned population; NPT, active non-position-tuned population. Blue and orange bars indicate time points or position bins with significant dSPN > iSPN or iSPN > dSPN differences, respectively. Gray bars indicate significant differences between population contrast traces in K-M. Shaded regions in all line plots are the 95% confidence intervals of the model coefficients from the linear mixed-effects model. Significance bars indicate p < 0.05, t-tests on model coefficients, Bonferroni corrected for multiple comparisons.

**Extended Data Fig. 8:**
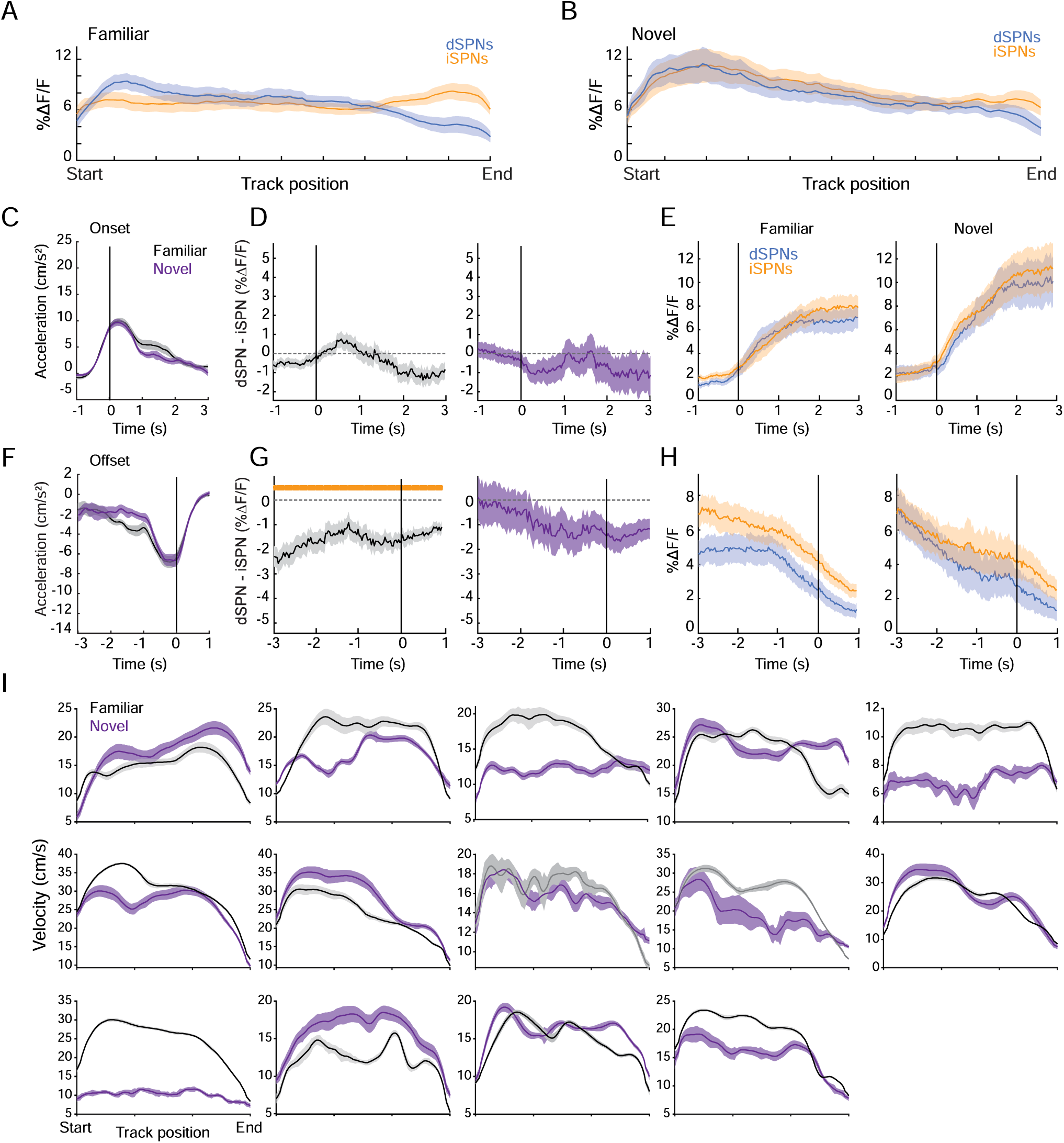
dSPN/iSPN activity during familiar and novel track traversal and low-amplitude locomotor transitions. **A**, Mean dSPN and iSPN population activity binned by track position during familiar track traversal (n = 14 mice, 14 sessions; 520 dSPNs, 836 iSPNs). B, Mean dSPN and iSPN population activity binned by track position during initial exposure to a novel track (n = 14 mice, 14 sessions; 592 dSPNs, 1050 iSPNs). C, Mean acceleration aligned to low-amplitude locomotor onset events during familiar and novel track traversal (familiar: n = 13 mice, 13 sessions; novel: n = 12 mice, 12 sessions). D, Mean dSPN-iSPN activity difference aligned to low-amplitude locomotor onset events during familiar and novel track traversal. E, Mean dSPN and iSPN population activity aligned to low-amplitude locomotor onset events during familiar and novel track traversal. F, Mean acceleration aligned to low-amplitude locomotor offset events during familiar and novel track traversal. G, Mean dSPN-iSPN activity difference aligned to low-amplitude locomotor offset events during familiar and novel track traversal. H, Mean dSPN and iSPN population activity aligned to low-amplitude locomotor offset events during familiar and novel track traversal. I, Velocity profiles for familiar and novel track traversal in individual mice. Blue and orange bars indicate time points or position bins with significant dSPN > iSPN or iSPN > dSPN differences, respectively. Shaded regions in all line plots are the 95% confidence intervals of the model coefficients from the linear mixed-effects model. Significance bars indicate p < 0.05, t-tests on model coefficients, Bonferroni corrected for multiple comparisons.

**Extended Data Fig. 9:**
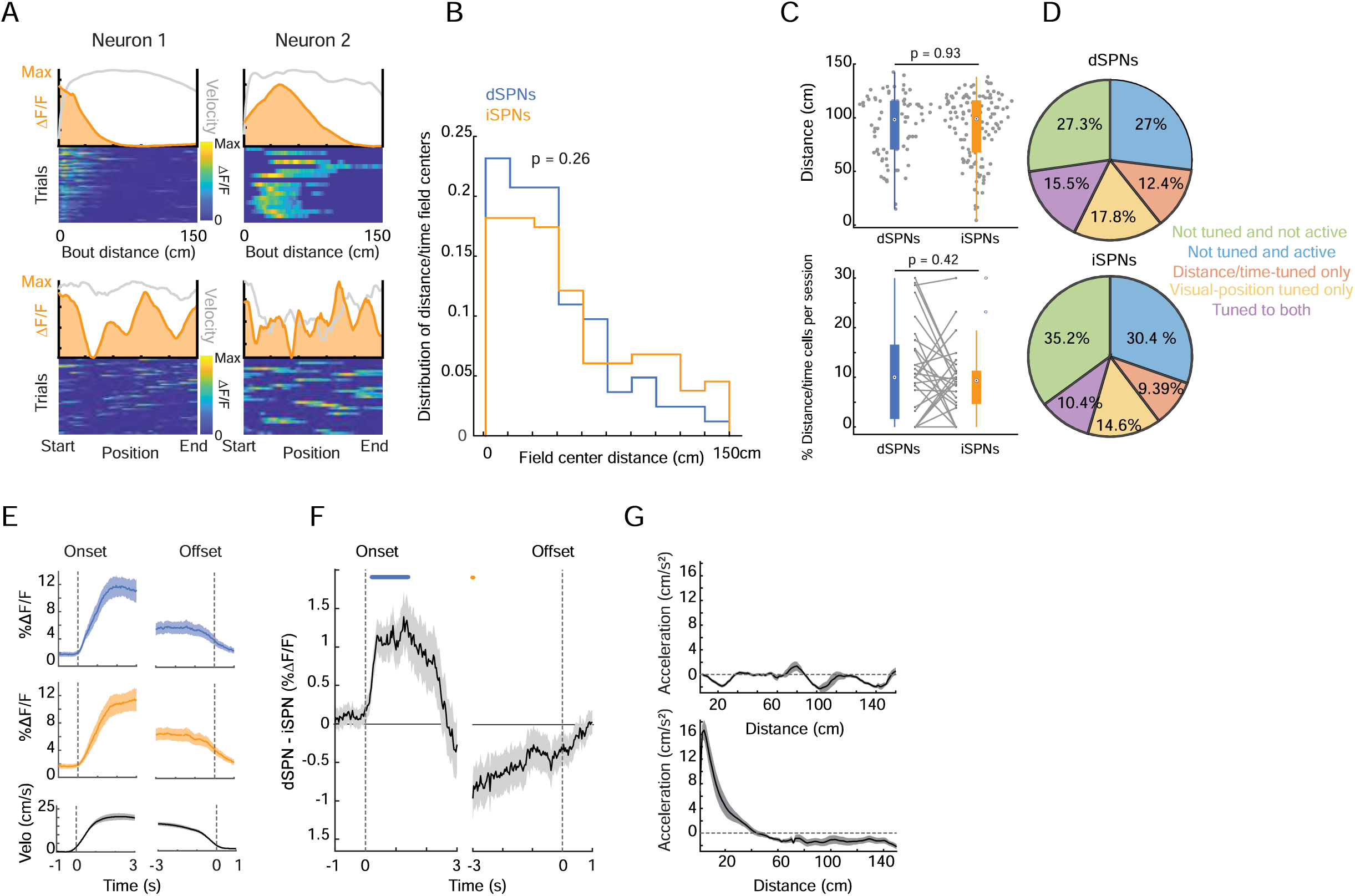
Additional characterization of distance/time and visual-position tuning in the infinite track task. **A**, Representative distance/time-tuned iSPNs shown with activity binned by bout distance/time or by visual position, corresponding to the example format in Fig. 5D. Line plots show mean ΔF/F with overlaid velocity in gray, and heatmaps show single-trial ΔF/F binned by the indicated variable. B_r_ Distribution of distance/time field centers for dSPNs and iSPNs (n = 82 dSPNs, 132 iSPNs; p = 0.26, Kolmogorov-Smirnov test). C, Quantification of distance/time tuning in dSPNs and iSPNs. Top, distance/time field-center location for tuned dSPNs and iSPNs. Bottom, percentage of distance/time-tuned cells per session for dSPNs and iSPNs (n = 28 sessions). Points indicate neurons or sessions as appropriate; paired lines indicate session-matched comparisons. P values indicate dSPN versus iSPN comparisons (paired, two-sided Wilcoxon signed-rank tests). D, Proportion of dSPNs and iSPNs classified as not tuned and not active, not tuned and active, distance/time tuned only, visual-position tuned only, or tuned to both distance/time and visual position. E, Mean dSPN activity, iSPN activity, and velocity aligned to locomotor onset and offset in the infinite track task (n = 7 mice, 26 sessions). F, Mean dSPN-iSPN activity difference aligned to locomotor onset and offset in the infinite track task. Positive values indicate greater dSPN activity and negative values indicate greater iSPN activity. G, Mean acceleration binned by visual position within the repeating track segment (top) or by bout distance from locomotor onset (bottom). Blue and orange bars indicate time points with significant dSPN > iSPN or iSPN > dSPN differences, respectively. Shaded regions in all line plots are the 95% confidence intervals of the model coefficients from the linear mixed-effects model. Significance bars indicate p < 0.05, t-tests on model coefficients, Bonferroni corrected for multiple comparisons.

